# Helices on interdomain interface couple catalysis in the ATPPase domain with allostery in *Plasmodium falciparum* GMP synthetase

**DOI:** 10.1101/808618

**Authors:** Santosh Shivakumaraswamy, Nivedita Pandey, Lionel Ballut, Sébastien Violot, Nushin Aghajari, Hemalatha Balaram

## Abstract

GMP synthetase catalyzes the substitution of the C2 oxo-group of the purine base in XMP with an amino-group generating GMP, the last step in the biosynthesis of GMP. This reaction involves a series of catalytic events that include hydrolysis of Gln generating ammonia in the glutamine amidotransferase (GATase) domain, activation of XMP to adenyl-XMP intermediate in the ATP pyrophosphatase (ATPPase) domain and reaction of ammonia with the intermediate to generate GMP. Inherent to the functioning of GMP synthetases is bidirectional domain crosstalk, which leads to allosteric activation of the GATase domain by substrates binding to the ATPPase domain, synchronization of the two catalytic events and tunnelling of ammonia from the GATase to the ATPPase domain. Herein, we have taken recourse to the analysis of structures of GMP synthetases, site-directed mutagenesis and, steady-state and transient kinetic assays on the *Plasmodium falciparum* enzyme to decipher the molecular basis of catalysis in the ATPPase domain and domain crosstalk. The results map the residues critical for catalysis in the ATPPase domain to the helices α11 and α12 that are located at the interdomain interface, and the lid-loop that follows α11. This apart, perturbing interdomain interactions involving residues on α11 and α12 impairs GATase activation. These results imply that this arrangement of helices at the domain interface with residues that play roles in ATPPase catalysis as well as domain crosstalk enables coupling ATPPase catalysis with GATase activation. Overall, the study enhances our understanding of GMP synthetases, which are drug targets in many infectious pathogens.

## Introduction

Glutamine amidotransferases are enzymes that operate in different biosynthetic pathways where they catalyze the substitution of an oxo group with an amino group^1^. Guanosine monophosphate synthetase (GMPS), a glutamine amidotransferase is a two domain (or two subunit) enzyme with modular catalytic units that catalyzes the substitution of the C2 oxo group of XMP with an amino group, producing GMP. This constitutes the last step in the *de novo* production of GMP. This conversion of XMP to GMP involves the coordinated functioning of the glutamine amidotransferase (GATase) domain that catalyzes the hydrolysis of the amide side chain of glutamine (Gln) producing ammonia and the ATP pyrophosphatase (ATPPase) domain that catalyzes the synthesis of adenyl-XMP intermediate from ATP.Mg^2+^ and XMP (Scheme 1)^1,2^. The ammonia generated in the GATase domain is tunnelled to the ATPPase domain where it attacks the adenyl-XMP intermediate generating GMP. *In vitro*, GMP synthetases can utilize externally available ammonia, although this is not physiologically relevant as at cellular pH, ammonia would be present largely as ammonium ion, which cannot serve as a nucleophile required for the reaction^3^. The GATase domain is allosterically activated by the binding of substrates to the ATPPase domain, thus ensuring that Gln hydrolysis occurs only when the ATPPase domain is primed to receive ammonia^4–6^. Further, the GATase and the ATPPase domains synchronize their reaction rates to avoid any futile utilization of Gln and ATP^7^. In some archaea, the GATase and the ATPPase domains are expressed separately (two-subunit type GMPS), and the two subunits interact during catalysis^8,9^. However, in most organisms, including archaea, the GATase domain (N-terminus) is connected to the ATPPase domain through a linker to form the two-domain type GMPS. Since GMP is involved in critical cellular functions, a functional GMPS is indispensable for the growth and survival of many organisms including infectious pathogens where the enzyme is pursued as drug targets^6,10–13^.

**Scheme 1.**
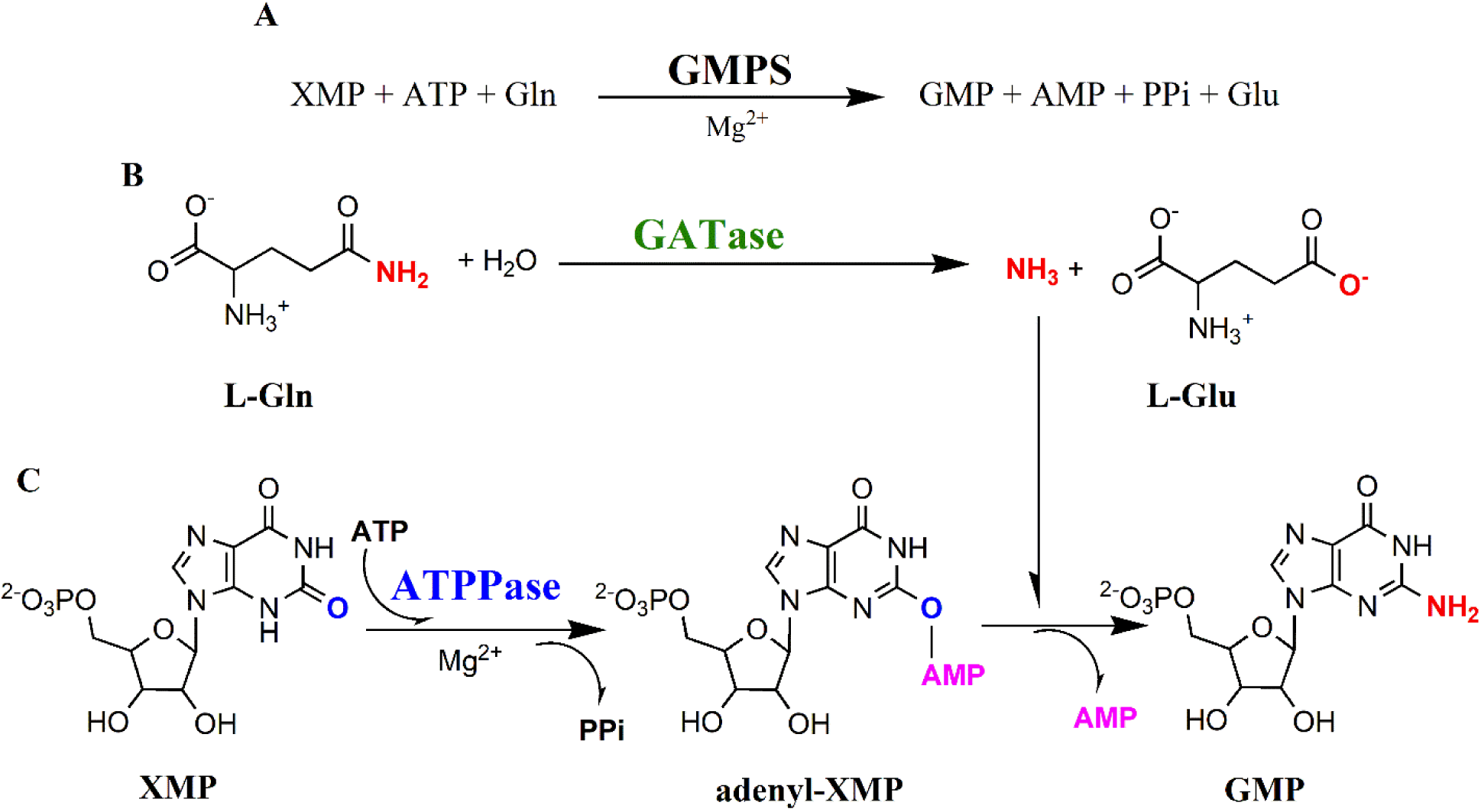
Reaction catalyzed by GMPS (A), GATase domain (B) and ATPPase domain (C).

The causative agent of cerebral malaria, *Plasmodium falciparum*, expresses a two-domain type GMPS (PfGMPS)^14^. Unlike other GMP synthetases, PfGMPS is unique in displaying basal GATase activity in the absence of ATPPase substrates^15^. However, binding of substrates to the ATPPase domain upregulates GATase activity by 8-12-fold^15,16^. A comparison of the kinetic parameters of the GATase reaction in the absence and presence of ATPPase substrates shows that GATase activation is enabled by lowering the *K*_m(app)_ value for Gln by 360-fold^16^. Apart from information flow from ATPPase to GATase domain, the inhibition of ammonia dependent GMP formation by Gln binding to a GATase inactive PfGMPS_C89A mutant suggests bidirectional interdomain crosstalk in PfGMPS^7^. This bidirectional domain crosstalk enables synchronization of the GATase and ATPPase reaction rates as evidenced by the 1:1 stoichiometry of the products of the two domains^7^.

The GATase domain in GMP synthetases has an α/β hydrolase fold and a Cys, His, Glu triad catalysing the hydrolysis of Gln^17^. The mechanism of substrate binding and catalysis in the GATase domain is well explored^3,5,18^, however, insights into the functioning of the ATPPase domain are lacking. Further, unlike in a majority of amidotransferases which possess either pre- formed or transient ammonia tunnels^19–21^, the crystal structures of GMP synthetases in apo and substrate-bound states have a solvent-exposed GATase active site and do not reveal a tunnel^16,17,22^. Regardless of the absence of a tunnel in the available structures, biochemical studies provide evidence for GATase activation and ammonia tunnelling in PfGMPS^7^. We recently reported the crystal structure of Gln bound PfGMPS_C89A mutant which revealed an 85° rotation of the GATase domain^16^. Upon reconstruction of the path traversed during domain rotation, a tunnel for the transport of ammonia is evident at an intermediate state of GATase rotation.

To understand the molecular basis of catalysis in the ATPPase domain and crosstalk with the GATase domain, we analyzed the structures of PfGMPS and other available GMP synthetases. Based on this analysis, eighteen site-directed mutants of PfGMPS were generated and subjected to enzyme assays to examine the activity of the individual domains, GATase activation and GMP formation. These studies have enabled the identification of residues that are involved in the catalysis of adenyl-XMP formation. In addition, our results highlight the involvement of residues on interface helices in the catalysis of adenyl-XMP formation and domain crosstalk. By tracing the connectivity of catalytic residues in the ATPPase domain with GATase residues, we present the molecular basis of active site synchronization. Our results on a dimer-interface mutant provide a functional rationale for the obligate dimeric nature of GMP synthetases from prokaryotes and parasitic protozoa, including PfGMPS, a feature distinct from most eukaryotic homologs that are monomers.

## Experimental Procedures

### Sequence and structure analysis

Protein sequences were retrieved from GenBank and UniProt sequence databases^23,24^. The sequence alignment was performed using the MAFFT web server^25^, and the figure was generated using the weblogo server^26^. The coordinates of the crystal structures of GMP synthetases were retrieved from RCSB Protein Data Bank^27^. The PDB IDs of the structures of GMPS analyzed are 1GPM (*Escherichia coli*), 3TQI (*Coxiella burnetii*), 2YWB and 2YWC (*Thermus thermophilus*), 5TW7 (*Neisseria gonorrhoeae*), 2VXO (*Homo sapiens*), 3UOW, 4WIM and 4WIO (*Plasmodium falciparum*). PyMOL was used to visualize and superpose structures, and generate figures^28^. H-bonds were identified using a distance cut-off of 3.2 Å between the donor and acceptor atoms. The negatively charged Asp, Glu residues were inferred to be forming a salt bridge with the positively charged Arg, Lys or His if the side chain carboxyl oxygen atoms of either Asp or Glu were within 4 Å of the side chain nitrogen atoms of Arg, His or Lys^29,30^. The atoms in the positively charged residues were Nζ in Lys; Nε, Nη1, or Nη2 in Arg, and Nδ1 or Nε2 in His. Every chain in a structure was analyzed independently. The disordered side chains were modelled using Swiss-PDB Viewer^31^. The modelled residues were then energy minimized using GROMOS96 force field implementation of Swiss-PDB Viewer, leaving the rest of the structure in the original conformation.

### Site directed mutagenesis

Site-directed mutants of PfGMPS (UniProt Q8IJR9, EC 6.3.5.2) were generated using the pET_PfGMPS expression construct^7^ as a template. The mutants were constructed using the protocol outlined in Quikchange II site-directed mutagenesis kit (Agilent technologies) with minor modifications. The sequences of the primers used in the PCRs are in Table S1. Phusion DNA polymerase (Thermo Scientific) was used for the PCRs, and the products were treated with 20 units of DpnI (New England Biolabs) at 37 °C overnight to digest the methylated template DNA. *E. coli* XL1-Blue competent cells were transformed with the PCR products, and the presence of mutation as well as the absence of errors in the open reading frame was confirmed by sequencing.

### Over-expression and purification of PfGMPS wildtype and mutants

The expression and purification of PfGMPS wildtype (WT) and mutants were performed as described earlier^16^. The WT was expressed using the pQE30_PfGMPS construct^15^ in a GMPS knockout *E. coli* strain AT2465 (*λ*^−^*, e14*−*, guaA21, relA1, spoT1* and *thiE1*) and the mutants were expressed in *E. coli* BL21-CodonPlus (DE3)-RIL cells transformed with pET_PfGMPS construct. The cells were grown in 4 l of terrific broth supplemented with 100 μg ml^-1^ ampicillin (WT) or 100 μg ml^-1^ ampicillin and 34 μg ml^-1^ chloramphenicol (mutants). Protein expression was induced by adding 0.5 mM isopropyl-β-D-thiogalactoside after the OD_600_ reached 0.8. The cells were harvested after 18 h of further incubation at 27 °C (WT) or 16 °C (mutants) by centrifugation. The cell pellet was resuspended in lysis buffer (50 mM Tris-HCl, pH 7.4, 10 % (v/v) glycerol, 2 mM β-mercaptoethanol) and stored at -80 °C.

The frozen cells were thawed on ice and supplemented with 2 mM β-mercaptoethanol and 1 mM phenylmethanesulfonyl fluoride. The cells were disrupted by sonication and the lysate was centrifuged at 30500 g, 4 °C for 45 min. The supernatant was loaded onto 2 ml HisTrap HP Ni-NTA column (GE Healthcare) connected to ÄKTA Basic HPLC (GE Healthcare). The column was pre-equilibrated with lysis buffer supplemented with 0.1 mM phenylmethanesulfonyl fluoride. The column with the bound protein was washed with step gradients of increasing imidazole concentration in lysis buffer and the bound proteins were eluted by ramping the imidazole concentration to 500 mM. The eluted fractions were examined by 12% (w/v) sodium dodecyl sulfate-polyacrylamide gel electrophoresis. The fractions containing PfGMPS in high amounts were pooled and concentrated using Amicon Ultra-15 centrifugal filters with 30 kDa cutoff membrane (Merck Millipore). The concentrated protein solution (about 2.5 ml) was injected into HiLoad 16/600 Superdex 200pg column (GE Healthcare) which was equilibrated with a buffer containing 20 mM Tris-HCl, pH 7.4, 10% (v/v) glycerol, 1 mM EDTA and 2 mM DTT. The flow rate employed was 0.3 ml min^-1^ and the eluted fractions were examined by 12 % (w/v) sodium dodecyl sulfate-polyacrylamide gel electrophoresis. The fractions containing pure protein were pooled, concentrated using centrifugal filters, flash-frozen in liquid nitrogen and stored in -80 °C freezer. The concentration of the protein was estimated using Bradford’s assay^32^.

### Circular dichroism measurements

Far-UV circular dichroism spectra were recorded using a J-810 spectropolarimeter (Jasco) using a quartz cell with 1 mm path length. The spectra were recorded from 200 to 260 nm with a bandwidth of 1 nm and a scan speed of 20 nm min^-1^. Each spectrum is an average of three scans. The concentration of the protein was 5 µM in a buffer containing 6.7 mM Tris-HCl, pH 7.4, 3.3 % (v/v) glycerol, 0.3 mM EDTA and 0.6 mM DTT.

### Analytical size-exclusion chromatography

The oligomeric state of the protein was determined using analytical size-exclusion chromatography. An HR 10/30 column packed with Sephadex 300 matrix attached to an ÄKTA basic HPLC system equipped with a UV-900 detector was employed. The column was equilibrated with a buffer containing 50 mM Tris-HCl, pH 7.4, 100 mM KCl and calibrated with the protein standards β-amylase, alcohol dehydrogenase, bovine serum albumin, carbonic anhydrase and cytochrome c (Sigma-Aldrich). 100 µl of the protein sample of concentration of 1 mg ml^-1^ was injected into the column individually and eluted at a flow rate of 0.5 ml min^-1^. The eluted proteins were detected by monitoring absorbance at 220 nm.

## Enzyme assays

### Assay for measuring GMP formation

Gln and NH_4_Cl dependent GMP synthesis were monitored at 25 °C using a continuous spectrophotometric assay as reported previously^15^. The reduction in absorbance due to the conversion of XMP to GMP was monitored at 290 nm using a Hitachi U2010 spectrophotometer. A Δε value of 1500 M^-1^ cm^−1^ was used to calculate the concentration of the product formed^33^. The standard assay consisted of 90 mM Tris-HCl, pH 8.5, 150 µM XMP, 2 mM ATP, 5 mM Gln, 20 mM MgCl_2_, 0.1 mM EDTA and 0.1 mM DTT in a total reaction volume of 250 µl. To measure NH_4_Cl dependent GMP formation, Gln in the reaction mix was replaced with 50 mM NH_4_Cl. The reaction was initiated by adding PfGMPS WT or mutants to a final concentration of 0.92 µM in the reaction mix. The steady-state kinetic parameters were obtained by measuring initial velocities over a range of substrate concentrations. When the concentration of one substrate was varied, the concentration of the other two substrates was fixed at saturating levels. XMP concentration was uniformly fixed at 0.15 mM for WT and mutants. ATP concentration was fixed at 2 mM for WT and mutants except for PfGMPS_K415L where it was at 5 mM. Fixed Gln concentration varied across the enzymes with WT and K415L at 5 mM, 10 mM for R25L, 15 mM for D413A, 30 mM for E213A and K376L, 40 mM for K24L and 50 mM for D412A. This was done to ensure that the concentration of the fixed substrate was saturating for each of the mutants. Initial velocity data measured at various substrate concentrations were fitted to the Michaelis–Menten equation [*v*=(*v*_max_×[S])/(*K*_m(app)_+[S])] by nonlinear regression using the software Prism 5 (GraphPad Software). A minimum of 10-14 substrate concentrations were used to determine the kinetic parameters. The kinetic parameters were determined twice and the results were used to calculate the mean and SEM. Statistical significance between WT and mutants was calculated using unpaired Student’s *t* test

### Assay for GATase activity

GATase activity was measured at 25 °C by estimating the concentration of glutamate formed using L-glutamate dehydrogenase (Sigma-Aldrich) as the coupling enzyme. In the coupled enzyme assay, the glutamate generated by the GATase domain is converted to α-ketoglutarate along with the concomitant reduction of NAD^+^ to NADH. Hence, Glu formed was monitored as an increase in absorbance at 340 nm in a continuous spectrophotometric assay. The concentration of Glu produced was estimated using a Δε value of 6220 M^-1^ cm^-1^. A 250 µl reaction mix contained 100 mM Tris-HCl, pH 8.5, 20 mM Gln, 150 µM XMP, 3 mM ATP, 20 mM MgCl_2_, 0.5 mM NAD^+^, 50 mM KCl, 0.1 mM EDTA and 0.1 mM DTT. The reaction mix was supplemented with an excess of L-glutamate dehydrogenase to ensure that the coupling enzyme was not limiting and the reaction was initiated by adding PfGMPS WT or mutants to a final concentration of 0.31 µM. The activity was expressed as a percentage of the activity of WT. The values from triplicate measurements were used to calculate the mean and SEM. This assay was also used to determine the binding affinities for ATP.Mg^2+^ and XMP in mutants that were inactive for GMP formation. However, mutants, which displayed < 40% GATase activation reported poorly in the coupled enzyme assay at low substrate concentrations and hence their binding affinities could not be measured.

### Stopped-flow assay

Stopped-flow assay to detect the formation of the adenyl-XMP intermediate was performed as reported previously^16^ using an SFM 300 stopped-flow device (BioLogic Science Instruments) equipped with MOS-200/M optics. The one-phase exponential decay in absorbance at 290 nm (A_290_) corresponding to the formation of adenyl-XMP intermediate was monitored using a 150 W Xe(Hg) lamp as a light source and TC-100/10F cuvette which provides a path length of 1 cm. The total flow rate was 8 ml s^-1^, which resulted in a dead time of 3.8 ms. The instrument was controlled using Biokine32 software, version 4.62, which allows accumulation of 8000 data points for each trace. Data were acquired every ms, and traces from three or more shots were averaged. The A_290_ of the averaged data was normalized to set the initial offset to zero. The normalized A_290_ was plotted against time and the resulting plots were fitted to equation 1:

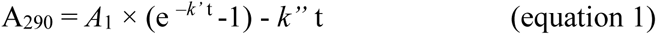

where *k*’ is the observed rate constant of the exponential phase corresponding to the formation of adenyl-XMP intermediate, *k*’’ is the rate constant of the steady-state phase and *A*_1_ is the amplitude of the exponential phase. A solution of 15 µM PfGMPS WT or mutants in 20 mM Tris-HCl, pH 7.4, 10 % (v/v) glycerol, 1 mM EDTA and 2 mM DTT was loaded in one syringe and the second syringe was loaded with a substrate mix containing 180 mM Tris-HCl, pH 8.5, 300 µM XMP, 4 mM ATP, 0.2 mM EDTA, 0.2 mM DTT and 40 mM MgCl_2_. The two solutions were mixed in a 1:1 ratio at a total flow rate of 8 ml s^-1^ and the A_290_ changes were recorded as a function of time. For each mutant, the exponential phase rate constant *k*’ was measured thrice and the *k*’ expressed as mean and SEM.

## Results

Examination of the available structures of GMP synthetases in conjunction with an alignment of GMPS sequences from diverse species enabled the identification of residues that could be involved in the catalysis of adenyl-XMP formation and interdomain crosstalk between the ATPPase and GATase domains. These observations were validated by generating eighteen site-directed mutants of PfGMPS and the positions of the mutated residues are shown in Figure 1A. The PfGMPS wild type (WT) and mutants were expressed in *Escherichia coli* and purified to greater than 95 % homogeneity (Figure S1). The absence of any significant difference in the far-UV circular dichroism spectra and elution times on analytical size-exclusion chromatography between the WT and mutants confirmed that the overall secondary structure and oligomeric state are retained (Figure S1). The mutants were then subjected to enzymatic assays to assign the role of the mutated residue to one of the three key steps in catalysis (Figure 1B) *viz*., adenyl-XMP formation, activation of GATase domain and GMP synthesis using either Gln or NH_4_Cl as nitrogen source with the former entailing ammonia tunnelling. The rate (*k*’) of adenyl-XMP formation was measured with ATP.Mg^2+^ and XMP as substrates using a stopped-flow spectrophotometer. The enhanced rate of Gln hydrolysis due to allosteric activation of the GATase domain by the binding of substrates to the ATPPase domain was measured using L-glutamate dehydrogenase as the coupling enzyme to detect the liberated glutamate. The same assay was also employed to determine the binding affinities (*K*_d_ values) of the inactive mutants for ATP.Mg^2+^ and XMP. The results of these experiments are discussed in the following sections.

**Figure 1.**
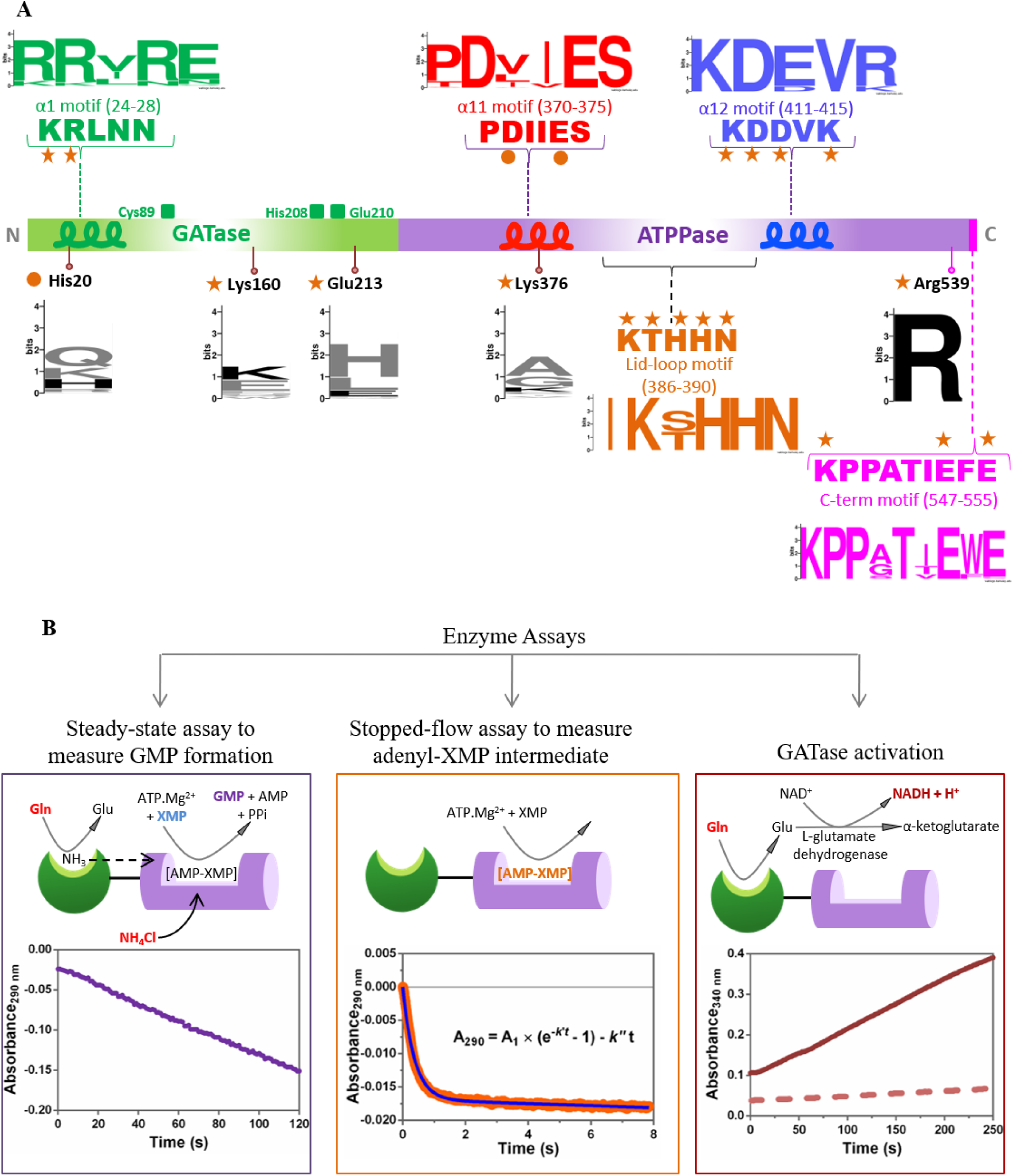
Overview of resides mutated and enzyme assays performed. (A) Schematic showing the location of the various conserved motifs that are examined in this study. The interface helices are shown as squiggles. The sequence of conserved motifs in PfGMPS, as well as the weblogo representation of the conserved motifs are shown. The residues investigated by site-directed mutagenesis in this and an earlier study^16^ are shown as stars and circles, respectively. The location of the GATase catalytic triad residues is shown as squares. (B) Scheme illustrating the enzymatic assays performed using PfGMPS WT and mutants. In the plot for adenyl-XMP formation, data is shown as points and the fit to the equation is shown as a line. The dashed line in the plot for GATase activation represents basal GATase activity and the solid line represents allosteric activation of GATase activity by substrates binding to the ATPPase domain.

### Catalysis in the ATPPase domain

Catalysis in the ATPPase domain involves the binding of substrates ATP.Mg^2+^, XMP and the subsequent formation of the adenyl-XMP intermediate through a phosphoryl transfer reaction. Here, the β-γ phosphoanhydride bond of ATP is cleaved and the AMP so generated is transferred to XMP^34,35^. The crystal structure of *E.coli* GMPS (EcGMPS) is complexed with AMP and PPi (PDB ID 1GPM)^17^ while the structures of human (PDB ID 2VXO)^22^, *T. thermophilus* (PDB ID 2YWC) and *P. falciparum* GMPS (PDB ID 3UOW) have a bound XMP. A 25-residue long lid-loop which connects the helix α11 with the strand β12 is disordered in all available structures except for human GMPS^22^ and PfGMPS_C89A/Gln (PDB ID 4WIO)^16^. The lid-loop is mapped away from the ATPPase active site in human GMPS; however, in PfGMPS_C89A/Gln, except for the eight intervening residues, the remaining residues of the lid-loop are mapped and positioned in the ATPPase active site (Figure 2A). Superposition of AMP-PPi and XMP from the structures of EcGMPS and PfGMPS, respectively, onto PfGMPS_C89A/Gln structure, revealed that residues that interact with ATP are primarily located in three regions *viz*., helices α11 and α12, the lid-loop and a conserved motif called the P-loop motif. Although residues from both the sub-domains interact with XMP, a majority of them are located on the C-terminal loop of the ATPPase domain. Considering that the role of the P-loop residues in catalyzing phosphoryl transfer reactions has been well studied^36,37^, we mutated the residues on helices α11 and α12, the lid-loop and the C-terminal loop to evaluate their role.

**Figure 2.**
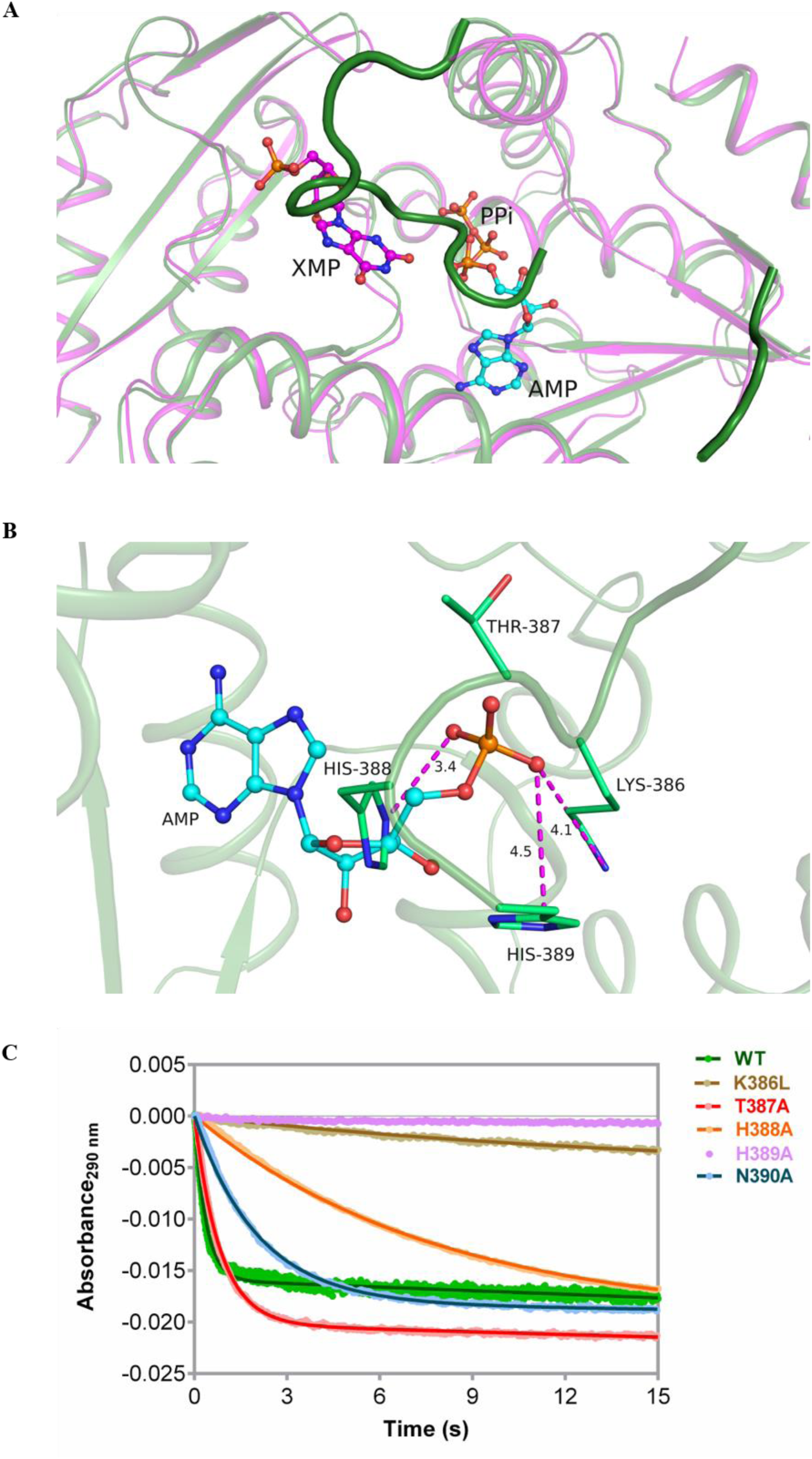
Role of lid-loop residues in adenyl-XMP formation. (A) Superposition of the ATPPase domain of XMP bound PfGMPS (magenta, PDB ID 3UOW) on the ATPPase domain of PfGMPS_C89A/Gln (green, PDB ID 4WIO). The lid-loop in PfGMPS_C89A/Gln is highlighted for reference. (B) Interactions of residues on the lid-loop with AMP in the structure of PfGMPS_C89A/Gln. Shown in dashed line are only salt bridge interactions. In both (A) and (B), AMP and PPi are superposed from EcGMPS (PDB ID 1GPM). Protein backbone is in ribbon and ligands are in ball and stick representation. (C) Stopped-flow progress curves for the formation of adenyl-XMP intermediate. Shown are absorbance values at 290 nm as a function of time (points) and fit of the data (solid lines) to one-phase exponential decay equation. The concentrations used were 7.5 µM enzyme, 2 mM ATP.Mg^2+^ and 0.15 mM XMP.

#### Invariant residues on the lid-loop

The lid-loop has an invariant motif IK(T/S)HHN (residues 385-390 in PfGMPS) (Figure 1A) and except for Asn390 which is disordered, other residues of the motif are in contact with AMP (superposed from EcGMPS) (Figure 2B). The residues KTHHN of the invariant motif were mutated one at a time and the purified mutants subjected to enzymatic assays. The Gln and NH_4_Cl dependent GMP formation by the PfGMPS mutants K386L, H388A and H389A were drastically impaired, ranging from 65-fold to 731-fold drop compared to WT (Table 1). Stopped- flow experiments revealed that the *k*’ for adenyl-XMP formation in PfGMPS_H388A was 24- fold lower than WT while the progress curves for K386L and H389A mutants did not display any drop in absorbance suggesting that these mutants were severely compromised in their ability to synthesize adenyl-XMP (Figures 2C and 3B).

**Figure 3.**
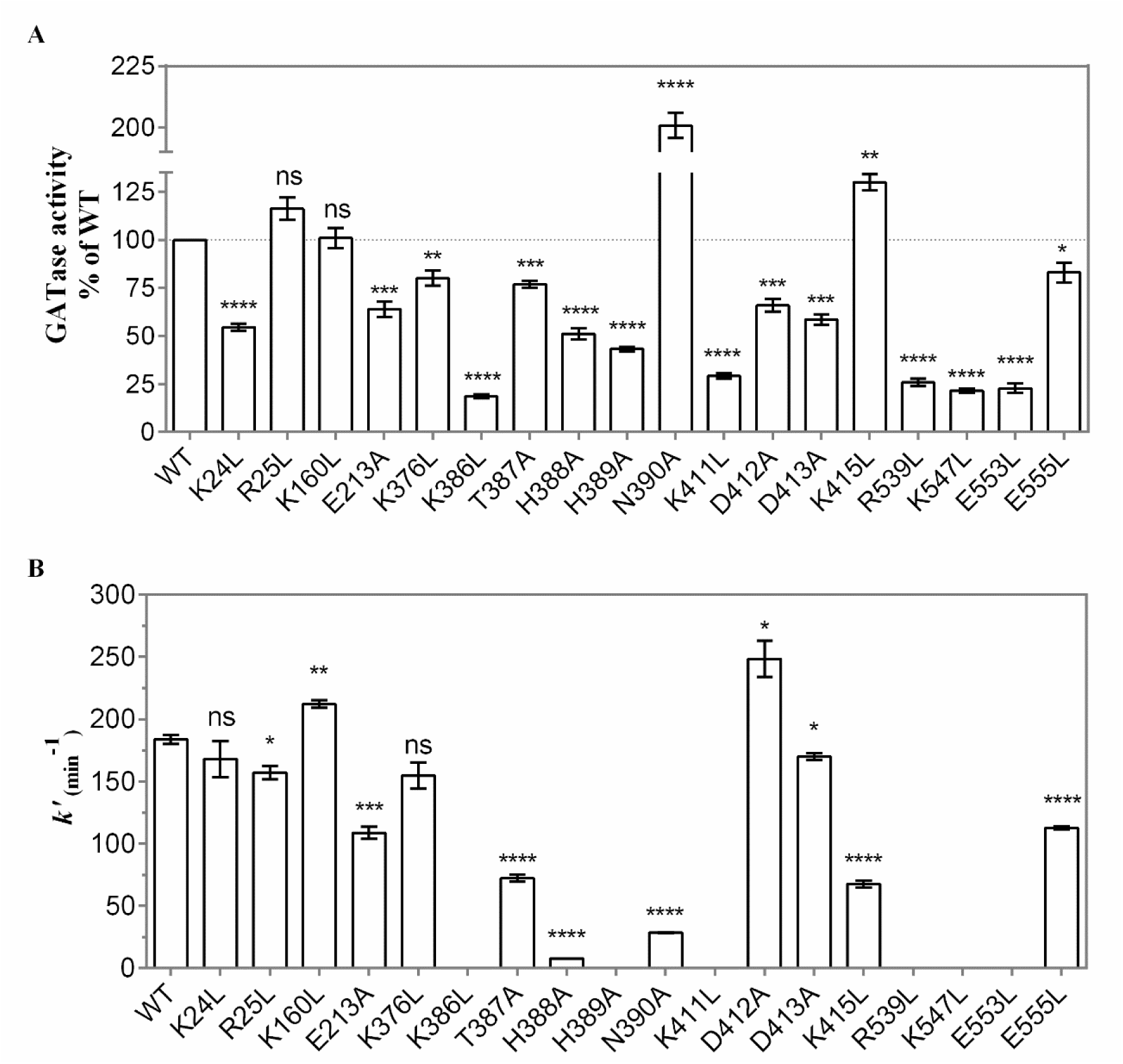
GATase activation and adenyl-XMP formation in PfGMPS WT and mutants. (A) Magnitude of GATase activation estimated using a coupled enzyme assay. GATase activity was measured using 20 mM Gln, 3 mM ATP.Mg^2+^ and 150 µM XMP. The specific activity values are normalized to the GATase activity of the WT. The error bars represent the SEM of data from three measurements. (B) Rate (*k*’) of adenyl-XMP formation measured using a stopped-flow spectrophotometer. The error bars represent the SEM of data from three independent measurements of *k*’. The concentration of the enzyme used was 7.5 µM and the concentration of ATP.Mg^2+^ and XMP were 2 mM and 0.15 mM, respectively. PfGMPS mutant is indicated by the residue, location in sequence and the residue to which it is mutated. In both A and B, the results of unpaired Student’s t test are indicated as follows, ns, not significant; *, p < 0.05; **, p < 0.01; ***, p < 0.001; ****, p < 0.0001.

**Table 1.**
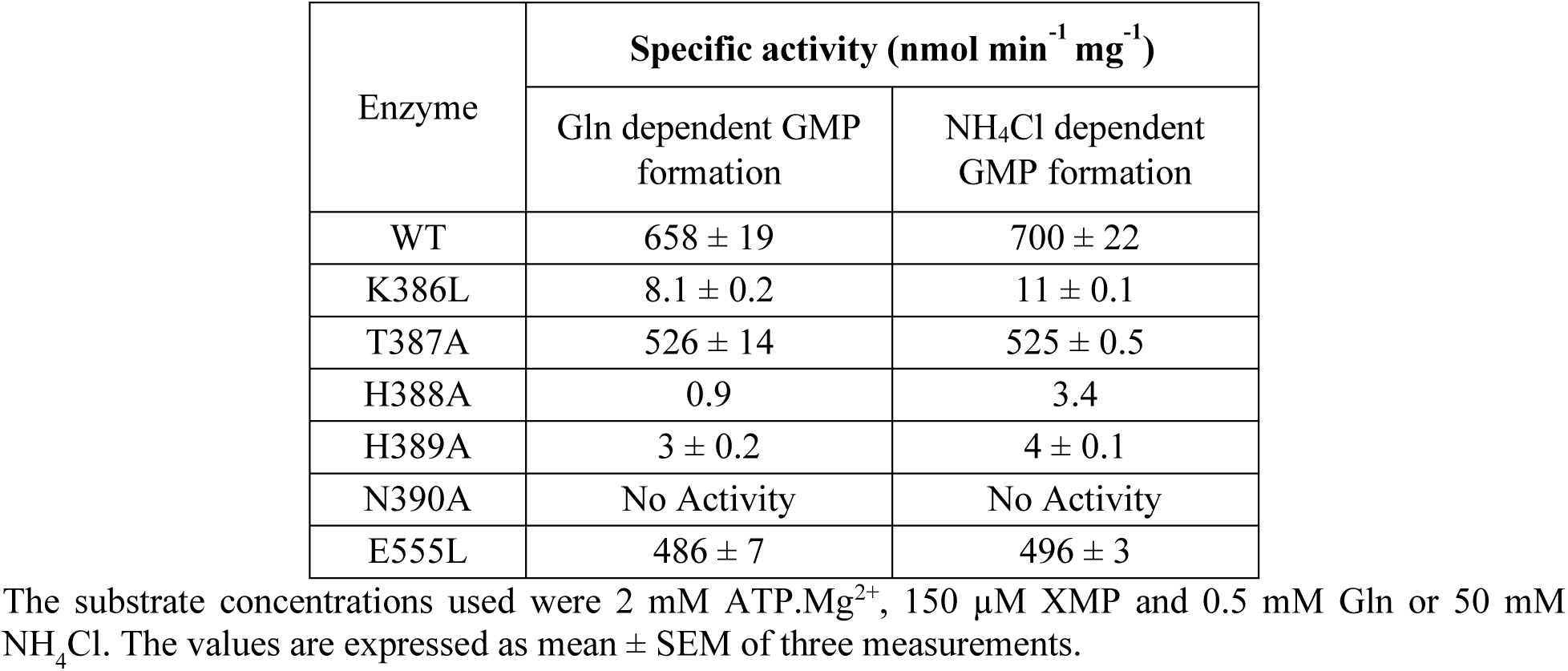
Gln and NH_4_Cl dependent GMP formation by PfGMPS WT, lid-loop and C-terminal loop mutants.

| Enzyme | Specific activity (nmol min<sup>-1</sup> mg<sup>-1</sup>) |  |
| --- | --- | --- |
|  | Gln dependent GMP formation | NH<sub>4</sub>Cl dependent GMP formation |
| WT | 658 ± 19 | 700 ± 22 |
| K386L | 8.1 ± 0.2 | 11 ± 0.1 |
| T387A | 526 ± 14 | 525 ± 0.5 |
| H388A | 0.9 | 3.4 |
| H389A | 3 ± 0.2 | 4 ± 0.1 |
| N390A | No Activity | No Activity |
| E555L | 486 ± 7 | 496 ± 3 |

The substrate concentrations used were 2 mM ATP.Mg<sup>2+</sup>, 150 µM XMP and 0.5 mM Gln or 50 mM NH<sub>4</sub>Cl. The values are expressed as mean ± SEM of three measurements.

To rule out impairment in substrate(s) binding being the cause for the lowered rate of adenyl-XMP/ GMP formation, we used GATase activation assay to measure affinities for ATP.Mg^2+^ and XMP. The levels of GATase activation in H388A and H389A mutants of PfGMPS were 51 and 43% of the WT (Figure 3A), respectively, while PfGMPS_K386L showed a massive drop to 19% that is only marginally greater than background basal (without binding of ATP.Mg^2+^ and XMP) glutaminase activity of PfGMPS. The *K*_d_ values for the complex of PfGMPS mutants H388A and H389A with ATP.Mg^2+^ and XMP are in the low micromolar range (Table S2) while the low level of activation in PfGMPS_K386L precluded *K*_d_ measurement. These results demonstrate that Lys386 is essential for substrate binding, whereas His388 and His389 play a catalytic role in the synthesis of adenyl-XMP intermediate.

Of the two other lid-loop mutants, PfGMPS_T387A displayed only a minor reduction in the specific activity for Gln and NH_4_Cl dependent GMP formation compared to WT while PfGMPS_N390A was completely inactive (Table 1). Both the mutants could synthesize adenyl-XMP intermediate albeit at reduced rates as the *k*^’^ for adenyl-XMP formation was 3-fold and 7- fold lower in the T387A and N390A mutants of PfGMPS, respectively (Figures 2C and 3B). The activation of the GATase domain in PfGMPS_T387A was reduced by 23% while it was enhanced to 200% of WT in PfGMPS_N390A (Figure 3A). These results establish that Thr387 is not important for PfGMPS catalysis while Asn390 plays an essential role in a step subsequent to adenyl-XMP formation as it can synthesize the adenyl-XMP intermediate but not the final product GMP.

#### Residues on inter-domain interface helices α11 and α12

In the crystal structures where the GATase domain is in 0° rotated state, the domain interface is composed of helix α1 (GATase) and the helices α11 and α12 (ATPPase) (discussed below). These helices harbour conserved motifs (Figure 1A), of which the side chains of some of the residues directed at the interface, engage in interdomain interactions. The side chains of the other residues *viz*., Asp371 (α11 motif) and Lys411 and Lys415 (α12 motif) are directed into the ATPPase catalytic pocket (Figure 4). The residue Asp371 has its side chain placed above the expected location of the adenyl-XMP bond. The side chain of Lys411 is involved in salt bridge like interactions with the phosphate of PPi that is distal to AMP (corresponding to γ-phosphate of ATP) while the side chain of Lys415 is devoid of contacts with the ligand or with other residues in PfGMPS structures (PDB IDs 3UOW, 4WIM).

**Figure 4.**
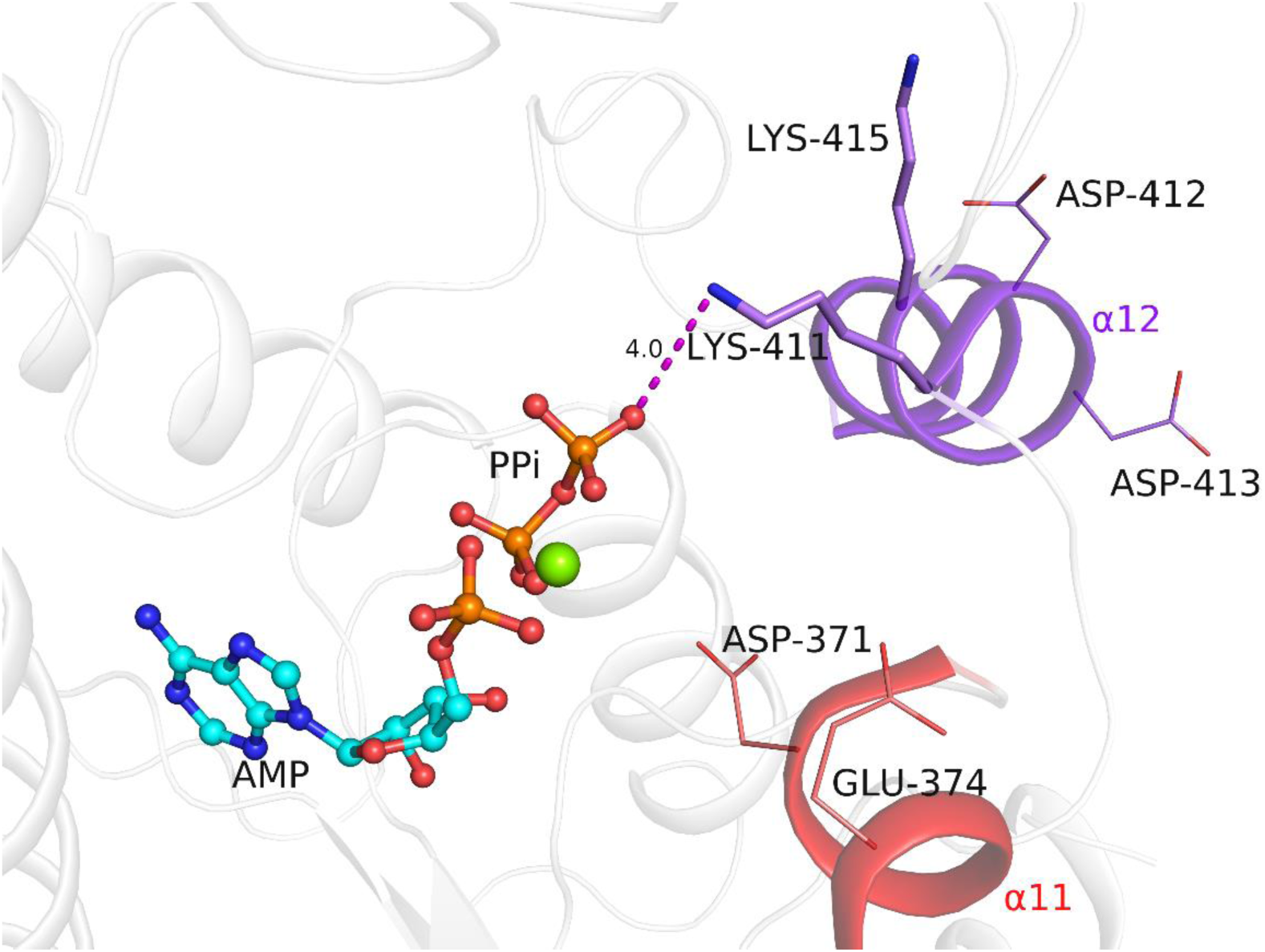
Interaction of Lys411 on the interface helix α11 with PPi. The ligands AMP, PPi (ball and stick) and Mg^2+^ (sphere) are superposed from the crystal structure of EcGMPS (PDB ID 1GPM). The helices α11 and α12 are coloured red and purple, respectively.

The almost complete impairment of adenyl-XMP formation upon mutation of Asp371 to Ala, reported in our previous study, indicates a key catalytic role for this residue on the interface helix α11^16^. Mutation of Lys411 on helix α12 to Leu resulted in a complete loss of Gln and NH_4_Cl dependent GMP formation, arising from the complete absence of adenyl-XMP formation in stopped-flow assays (Figure 3B). However, unlike in PfGMPS_D371A, where GATase activation was higher than WT^16^, the 70% drop seen in PfGMPS_K411L indicates that substrate(s) binding is highly compromised (Figure 3A). Mutation of Lys415 to Leu altered only the *K*_m(app)_ value for ATP that was 4.2-fold higher compared to WT (Table 2), and this is reflected in 63% lower *k*’ for adenyl-XMP formation (Figure 3B). These results highlight the role of residues on the interface helices in substrate binding (Lys411, Lys415) and adenyl-XMP formation (Asp371). It should be noted that although the side chain of Lys415 (α12 motif) is devoid of any interactions in PfGMPS structures, in most other GMPS structures, the side chain of this residue is involved in H-bonding interactions with main chain atoms of residues lining the ATPPase pocket.

**Table 2.** Steady-state kinetic parameters of interdomain interface mutants.

| Varied substrate |  | WT | K24L | R25L | K160L | E213A | K376L | K411L | D412A | D413A | K415L |
| --- | --- | --- | --- | --- | --- | --- | --- | --- | --- | --- | --- |
| Gln | $K_{m(app)}$ (mM) | 0.33<br>± 0.02 | 1.75<br>± 0.06 | 0.47<br>± 0.01 | 0.363<br>± 0.004 | 1.29<br>± 0.04 | 1.46<br>± 0.07 | Inactive | 2.61<br>± 0.22 | 1.19<br>± 0.10 | 0.25<br>± 0.01 |
|  |  |  | ** | * | ns | ** | ** |  | ** | * | ns |
| | $k_{cat}$ (s<sup>-1</sup>) | 0.766<br>± 0.005 | 0.296<br>± 0.007 | 0.777<br>± 0.031 | 0.832<br>± 0.007 | 0.794<br>± 0.026 | 0.804<br>± 0.022 | | 0.657<br>± 0.004 | 0.584<br>± 0.008 | 0.707<br>± 0.004 |
|  |  |  | *** | ns | * | ns | ns |  | ** | ** | * |
| | $k_{cat}/K_{m(app)}$ (s<sup>-1</sup> M<sup>-1</sup>) | 2339 | 169 | 1657 | 2292 | 614 | 553 | | 252 | 492 | 2780 |
| ATP | $K_{m(app)}$ (μM) | 119<br>± 2 | 67<br>± 3 | 117<br>± 4 | 148<br>± 3 | 162<br>± 2 | 122<br>± 0.1 | | 96<br>± 5 | 102<br>± 1 | 500<br>± 11 |
|  |  |  | ** | ns | * | ** | ns |  | ns | * | *** |
| | $k_{cat}$ (s<sup>-1</sup>) | 0.893<br>± 0.039 | 0.340<br>± 0.004 | 0.831<br>± 0.003 | 0.820<br>± 0.019 | 0.846<br>± 0.009 | 0.872<br>± 0.008 | | 0.724<br>± 0.022 | 0.598<br>± 0.028 | 0.733<br>± 0.018 |
| | $k_{cat}/K_{m(app)}$ (s<sup>-1</sup> M<sup>-1</sup>) | 7496 | 5112 | 7099 | 5553 | 5226 | 7161 | | 7547 | 5855 | 1465 |
| XMP | $K_{m(app)}$ (μM) | 7.1<br>± 1 | 20.6<br>± 1 | 6.3<br>± 0.4 | 12.7<br>± 0.6 | 13.1<br>± 1 | 12.1<br>± 0.6 | | 9.5<br>± 1.9 | 20<br>± 2.9 | 14.4<br>± 1.5 |
|  |  |  | * | ns | * | * | * |  | ns | ns | ns |
| | $k_{cat}$ (s<sup>-1</sup>) | 0.741<br>± 0.058 | 0.327<br>± 0.005 | 0.751<br>± 0.070 | 0.936<br>± 0.035 | 0.891<br>± 0.025 | 0.956<br>± 0.029 | | 0.633<br>± 0.058 | 0.532<br>± 0.011 | 0.744<br>± 0.059 |
| | $k_{cat}/K_{m(app)}$ (× 10<sup>3</sup> s<sup>-1</sup> M<sup>-1</sup>) | 104 | 15.9 | 119 | 73.8 | 68.2 | 79.2 | | 66.9 | 26.7 | 51.7 |
| NH<sub>4</sub>Cl | $K_{m(app)}$ (mM) | 25.6<br>± 2.8 | 20.3<br>± 2.1 | 13.1<br>± 1.1 | 23.6<br>± 0.6 | 12.3<br>± 0.4 | 36.2<br>± 2.2 | | 38.9<br>± 0.2 | 12.2<br>± 1.4 | 17<br>± 0.3 |
|  |  |  | ns | ns | ns | * | ns |  | * | * | ns |
| | $k_{cat}$ (s<sup>-1</sup>) | 1.081<br>± 0.107 | 0.465<br>± 0.011 | 0.939<br>± 0.004 | 1.240<br>± 0.027 | 0.708<br>± 0.002 | 0.792<br>± 0.073 | | 1.054<br>± 0.030 | 0.711<br>± 0.024 | 0.591<br>± 0.008 |
|  |  |  | * | ns | ns | ns | ns |  | ns | ns | * |
| | $k_{cat}/K_{m(app)}$ (s<sup>-1</sup> M<sup>-1</sup>) | 42 | 23 | 72 | 53 | 57 | 22 | | 27 | 58 | 35 |

The values are expressed as mean ± SEM of two independent experiments. The assay involved monitoring the decrease in absorbance at 290 nm arising from the conversion of XMP to GMP at 25 °C. The results of unpaired Student's t test are indicated as follows, ns, not significant; \*, p < 0.05; \*\*, p < 0.01; \*\*\*, p < 0.001; \*\*\*\*, p < 0.0001.

#### Residues on the C-terminal loop

The C-termini of the ATPPase domain in GMP synthetases comprises a 13-residue long loop (C-terminal loop) which bears the highly conserved signature motif KPPXTXE(F/W)X (residues 547-555 in PfGMPS) (Figure 1A). The ζ-amino group of Lys547 interacts with the phosphate of XMP (Figure 5A). The peptide bonds Lys547-Pro548-Pro549 are in the *cis* configuration, which, induces a turn in the polypeptide chain and enables the interaction of Glu553 side chain carboxyl group with ribose 2’ hydroxy group of XMP (Figure 5A). In addition, the α-carboxy of the terminal Glu555 residue forms a salt bridge with the guanidino group of Arg336 that in turn interacts with C6 oxygen and N7 of XMP (Figure 5A). Hence, residues of the C-terminal loop make direct and indirect contacts with XMP. The conformation of the loop is restrained by dimer interactions involving H-bonds between the Nη2 of Arg539’ (from the neighbouring chain) and backbone CO of both Thr551 and Glu553 (Figure 5B).

**Figure 5.**
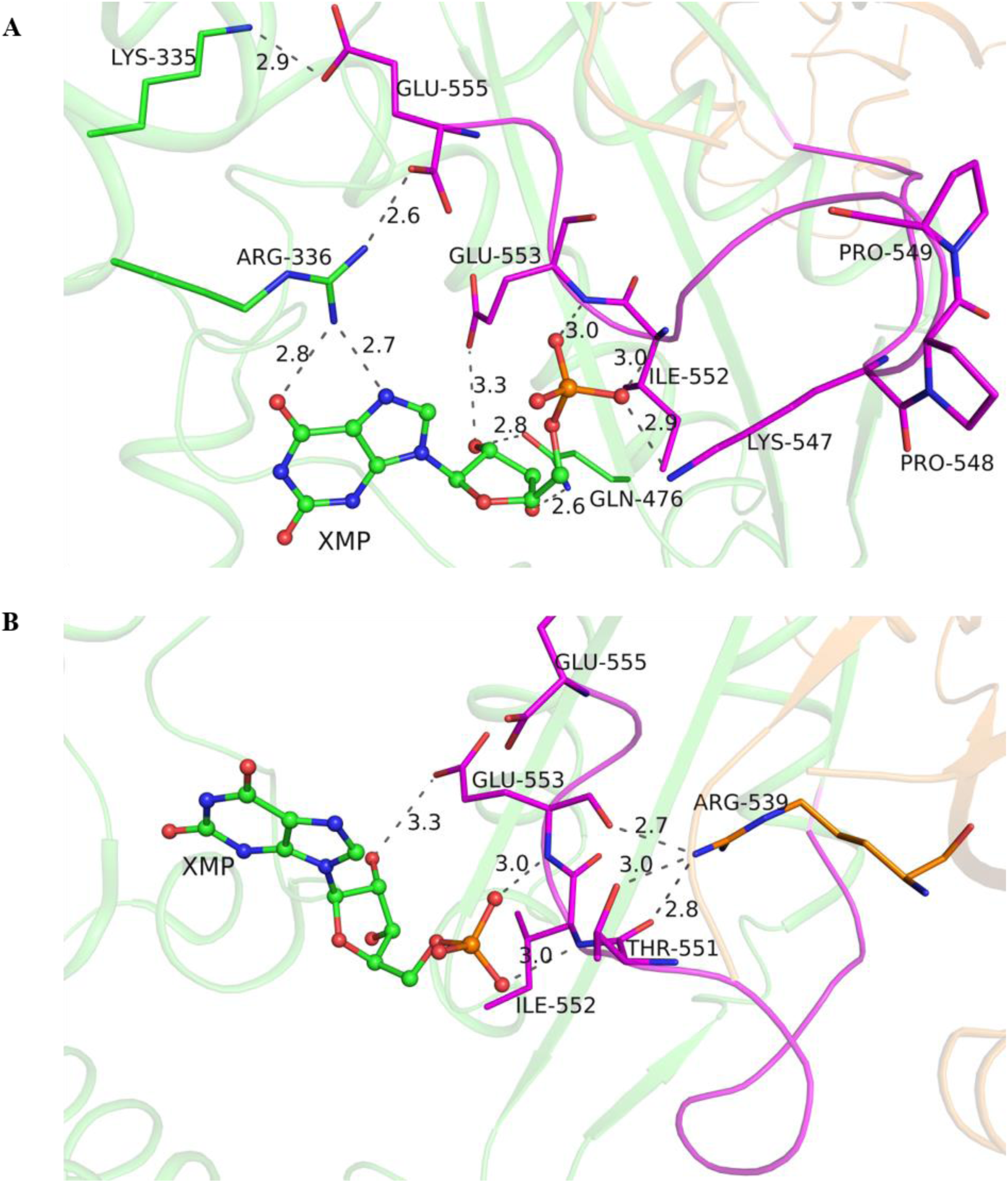
Interactions of the C-terminal loop in PfGMPS (PDB ID 3UOW). (A) Interactions with XMP. (B) Interdimer interactions of Arg539 from a neighbouring chain with residues on C-terminal loop. In both images, the protein backbone is in ribbon representation with chain A coloured green and chain B orange. The C-terminal loop of chain A is coloured magenta. XMP is shown as ball and stick and highlighted residues are shown as sticks.

We probed the role of C-terminal loop residues Lys547, Glu553, Glu555 as well as the importance of the interdimer interactions of Arg539 by mutating them to Leu. The K547L, E553L and R539L mutants of PfGMPS were inactive for Gln and NH_4_Cl dependent GMP formation on account of impaired adenyl-XMP formation (Figure 3B). GATase activation was reduced by >70% suggesting that these mutants are defective in binding substrates (Figure 3A). In contrast, the Gln and NH_4_Cl dependent GMP formation by the PfGMPS_E555L mutant was only marginally lower than WT (Table 1). Also, the changes in GATase activation were minor, and the *k*’ for adenyl-XMP formation was lower by only 39% (Figure 3). These results prove that the direct interactions of Lys547 and Glu553 and the indirect interactions of Arg539 are necessary for substrate binding and corroborate structural observations.

### Domain crosstalk

#### Interdomain interface in GMP synthetases

The available GMPS structures were examined to identify residues at the interdomain interface that could mediate domain crosstalk. The GATase domain is oriented with respect to the ATPPase domain in the same manner in all GMPS structures except in the PfGMPS_C89A/Gln structure where this domain is rotated by 85° (Figure 6A, C). We refer to the state of the domain rotation in PfGMPS_C89A/Gln complex as the 85° and the others as 0°. In structures where the GATase is in 0° rotated state, the interdomain interface is predominantly composed of the N-terminal helix, α1 of the GATase domain and helices α11 and α12 of the ATPPase domain (Figure 6A). The three interface helices harbour the conserved motifs (K/R)(K/R)XRE (α1 motif), DXXES (α11 motif) and KD(D/E)V(K/R) (α12 motif) (Figure 1A). The charged residues in the 4^th^ and 5^th^ positions of the α1 motif are replaced with Asn in sequences from *Plasmodium* species. Further, the conserved D/E and K/R substitutions in the α12 motif are limited to sequences from *Plasmodium*.

**Figure 6.**
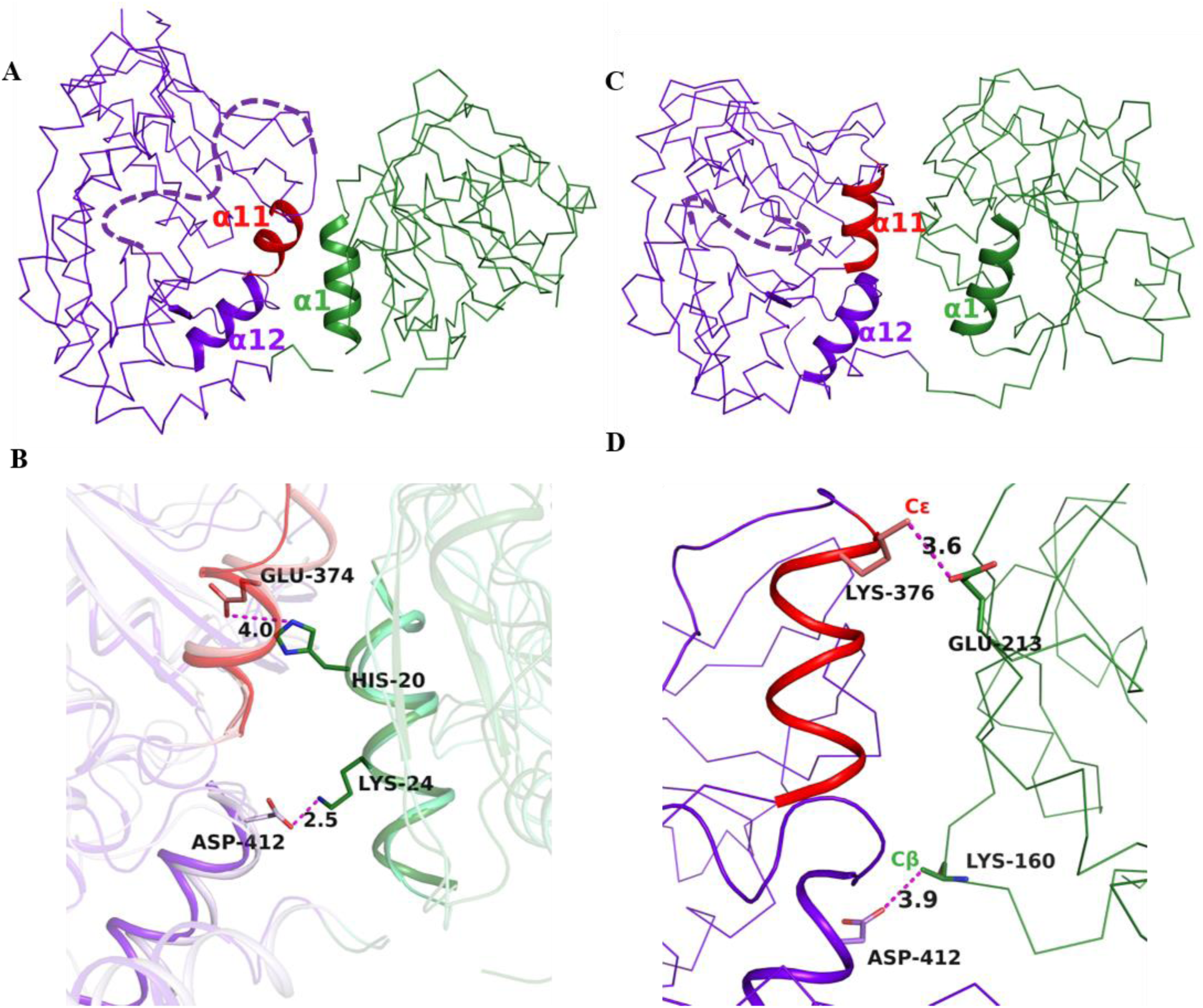
Interdomain interactions in PfGMPS. (A) Helices α1, α11 and α12 in the interdomain interface in the 0° structure of PfGMPS (PDB ID 3UOW). (B) Interdomain salt bridges in the 0° structures. The chain A in structure 3UOW is superposed on chain B of 4WIM (apo PfGMPS), which is shown in a lighter shade of colour. (C) Helices α11 and α12 in the interdomain interface in 85° GATase rotated PfGMPS structure (PDB ID 4WIO). (D) Interdomain salt bridges in the 85° rotated structure. In all the images, the GATase and ATPPase domains are coloured green and purple, respectively. The backbone is shown as line (A, C, D) and as ribbon (B). The disordered lid-loop is shown as a dashed line (A, C).

#### Interdomain interactions of the 0° interface

The α1-α12 interdomain interactions are largely conserved in all GMPS structures where the GATase is in the 0° rotated state. The first and the fourth residues of α1 motif ((K/R)(K/R)XRE) form two interdomain salt bridges with the second and third residues of the α12 motif (KD(D/E)V(K/R) (Table S3), respectively, though in *Neisseria gonorrhoeae* GMPS, the distance between the interacting pairs is above 4 Å. In the structures of PfGMPS (PDB ID 3UOW, 4WIM), the side chains of some of these residues are disordered. However, through modelling of missing side chains or structural superposition of protomers from two structures (3UOW and 4WIM), the atoms involved in salt bridge formation between Lys24 (α1) and Asp412/Asp413 (α12) come within a distance of ≤ 4 Å (Figure 6B and Table S3). The second salt bridge is absent in PfGMPS as the fourth residue of the α1 motif is an asparagine.

Unlike α12, the residues of α11 across the GMPS structures, show varying levels of disorder indicating conformational flexibility of this segment. Also, an interdomain salt bridge involving His20 preceding the α1 motif and Glu374 of α11 motif was observed by superposing two chains of PfGMPS (Figure 6B)^16^. His20 is not conserved across GMPS sequences (Figure 1A) and a similar salt bridge is observed only in *N. gonorrhoeae* GMPS where the histidyl residue is replaced by Arg (Table S3). Other interdomain salt bridges are enzyme-specific and the interacting residues are not conserved.

The significance of α1-α12 interdomain interactions was investigated in PfGMPS by mutating Lys24 of the α1 motif and Asp412 as well as Asp413 of the α12 motif. Compared to WT, the mutants K24L, D412A and D413A of PfGMPS displayed an increase in *K*_m(app)_ value for the substrate glutamine with PfGMPS_D412A showing the highest (7.9-fold) increase (Table 2). The *k*_cat_ for Gln dependent GMP formation was unaltered in the D412A and D413A mutants of PfGMPS, however, this parameter was lower by 60% in the K24L mutant compared to WT (Table 2). This reduction in *k*_cat_ value for PfGMPS_K24L was deemed significant by statistical test (*p* < 0.001). The reduction in *k*_cat_/*K*_m(app)_ for Gln dependent GMP formation in PfGMPS mutant K24L was the highest (13.9-fold), followed by D412A (9.3-fold) and D413A (4.8-fold). The levels of GATase activation were lower by 45%, 34% and 42% in the K24L, D412A and D413A mutants, respectively, while the *k*’ for adenyl-XMP formation did not vary significantly (Figure 3). Though PfGMPS_K24L is a GATase mutant, the *k*_cat_ for NH_4_Cl dependent GMP formation, a step that is independent of GATase activity, is reduced by half (Table 2). The second residue of the α1 motif, Arg25, interacts with the residues on the loop containing His208 and Glu210, which together with Cys89 constitute the GATase catalytic triad (Figure S2). Mutation of Arg25 to Leu however did not result in any significant change in kinetic properties, the level of GATase activation or the *k*’ for adenyl-XMP formation (Table 2 and Figure 3).

Unlike the mutation of lid-loop residues which results in complete inactivation, interface residues are distal to the GATase and ATPPase active site pockets and hence the observed changes are not drastic. However, the significant reduction in *k*_cat_/*K*_m(app)_ upon mutating Lys24, Asp412 and Asp413 suggests an essential role for these residues in mediating domain crosstalk. This is supported by the results of our previous study where we had probed the α1-α11 interactions. In a manner identical to Lys24, mutation of His20 on α1 results in a two-fold reduction in *k*_cat_ for both Gln and NH_4_Cl dependent GMP formation while the results of mutating the interacting partner, Glu374 on α11, suggested that this residue plays multiple roles including domain crosstalk and adenyl-XMP formation^16^.

#### Interdomain interactions of the 85° interface

In the PfGMPS_C89A/Gln structure (PDB ID 4WIO), helices α11 and α12 retain the conformation that is observed in the 0° structure, however, they interact with a different set of GATase residues on account of GATase rotation (Figure 6C). Although the side chains of many residues constituting this 85° interface are disordered, an examination of the structure shows that the side chains of the residue pairs Glu213/Lys376 and Lys160/Asp412 could be involved in forming interdomain salt bridges (Figure 6D). The GATase residues Glu213, Lys160 and the ATPPase residue Lys376 located following the α11 motif, are not conserved while Asp412 is a member of the invariant α12 motif (Figure 1A).

Mutation of Glu213 to Ala as well as Lys376 to Leu results in an increase in the *K*_m(app)_ value for Gln by 3.9 and 4.4-fold, respectively (Table 2) whereas the changes in *k*_cat_ are minor. In PfGMPS_E213A, the levels of GATase activation and the *k*’ for adenyl-XMP formation are reduced by 36% and 41%, respectively, while the changes for the K376L mutant are not significant (Figure 3). Mutation of Lys160 to Leu did not affect the kinetic parameters (Table 2), the levels of GATase activation or the *k’* for adenyl-XMP formation (Figure 4). However, as discussed above, mutation of Asp412, which is also involved in interdomain salt bridges in the 0° interface, impairs domain crosstalk. The increase in the *K*_m(app)_ value for Gln upon mutating Glu213 and Lys376, suggests a role for even the 85° interface residues in domain crosstalk.

## Discussion

The incorporation of nitrogen into XMP catalyzed by GMP synthetases requires the production of a reactive nitrogen atom (with its lone pair of electrons) in the GATase domain, activation of the substrate XMP in the ATPPase domain and the interdomain transfer of nitrogen (ammonia). In addition, GMP synthetases are characterized by allostery as events in the ATPPase domain upregulate Gln hydrolysis in the GATase domain. Also, the catalytic events in the two domains are synchronized. To identify the residues involved in these processes, we resorted to structural analysis, site-directed mutagenesis and enzyme assays. These investigations have helped us map the ATPPase active site as well as the residues involved in domain crosstalk.

### Mapping functional residues in the ATPPase catalytic pocket

Our studies have revealed that residues that are critical for the functioning of the ATPPase domain are located on the interface helices α11 and α12, lid-loop and C-terminal loop. Whereas Asp371 on α11 is essential for adenyl-XMP formation^16^, Lys411 and Lys415 as shown in this study, are involved in the binding of ATP. Interestingly, helices α11 and α12 also harbour residues whose side chains contact the GATase domain implicating a dual function of catalysis in the ATPPase domain and interdomain crosstalk (see below).

The lid-loop subsequent to helix α11 contains the invariant IK(T/S)HHN motif. The residue Lys386 participates in ATP binding, whereas His388 and His389 are involved in catalyzing the formation of the adenyl-XMP intermediate. The increase in electrophilicity of C2 of XMP enabled through the formation of the unstable adenyl-XMP intermediate facilitates attack by ammonia and thereby the replacement of the C2 oxygen with nitrogen. Transfer of AMP from ATP to XMP is an adenylation reaction that is facilitated by the ATP α-phosphorous undergoing geometric as well as electrostatic changes involving the development of strong negative charges on the non-bridging oxygen atoms in the transition state^38,39^. Since stabilization of transition state underlies enzymatic rate enhancements, all phosphoryl transfer enzymes, including adenylating enzymes, employ positively charged groups to interact with and neutralize the negative charges on the non-bridging oxygen atoms^40^. Our results suggest a dominant role for His389 and also a possible function for His388 in the stabilization of the transition state as both H388/389A mutants of PfGMPS bind ATP.Mg^2+^ and XMP, but are highly or completely impaired in adenyl-XMP formation. Crystal structures also lend support to the catalytic role of His388 and His389 as these two residues along with Lys386 are the only ones with positively charged side chains lying within an interacting distance of ATP α-phosphate.

The lid-loop is disordered in most structures of GMPS suggesting that it is conformationally dynamic. This is also seen in the adenylating enzyme tyrosyl-tRNA synthetase, wherein the loop containing the conserved KSMKS motif changes its conformation from an open to a closed-form upon binding ATP. This closure results in a movement of the ζ-amino group of Lys235 (KSMKS motif) close to the ATP α-phosphate thereby resulting in catalysis^41,42^. The results with the PfGMPS lid-loop mutants suggest that even in GMP synthetases, binding of ATP.Mg^2+^ and XMP induces conformational changes in the lid-loop culminating in the repositioning of the IK(T/S)HHN motif residues in proximity to ATP α-phosphate, facilitating the synthesis of adenyl-XMP intermediate through residues His388 and His389 as well as the formation of GMP through interactions of Asn390. This design of placing catalytic residues on a conformationally dynamic lid-loop has enormous functional implications as it ensures that the labile adenyl-XMP intermediate is synthesized only after a closed catalytic pocket is established. The sealed pocket protects the adenyl-XMP intermediate from attack by water and also enables ammonia channelling.

The C-terminus of the ATPPase domain mediates dimerization in PfGMPS and contributes many residues that interact with the substrate XMP. Our experiments suggest that residues Lys547 and Glu553 on the C-terminal loop are critical for binding XMP. Moreover, the interactions of Arg539’ from the neighbouring protomer with the C-terminal loop facilitates substrate binding by maintaining the correct conformation of the C-terminal loop^16^. The contribution of residues from the neighbouring protomer in substrate binding sheds light on the essentiality of dimerization in PfGMPS and GMP synthetases from prokaryotes and parasitic protozoa^6,13,15^. Human GMPS and GMP synthetases from higher eukaryotes possess a 150- residue insertion that enables them to function as monomers^43^. These results highlight a significant difference between the active sites of the ATPPase domain of GMPS from pathogens and the human host, which could be exploited in the design of specific inhibitors.

### Mechanism of allostery and domain crosstalk in GMP synthetases

The GATase domain is connected to the ATPPase domain through a flexible linker which permits an 85° rotation. In structures of two-domain type GMPS where the GATase is in 0° state, the interdomain interactions mediated by GATase helix α1 and ATPPase helices α11 and α12 are conserved. Even upon 85° rotation, helices α11 and α12 remain part of the interface, suggesting their central role in domain crosstalk. A total of nine residues in the 0° and 85° interfaces were mutated and the enzymatic assays implicate the interface residues in domain crosstalk.

As reported earlier, GATase activation induced by the binding of ligands to the ATPPase domain is characterized by a reduction in the *K*_m(app)_ value for Gln, arising from the formation of a high-affinity binding pocket^16^. Disruption of the interdomain salt bridges between Lys24 (helix α1) and Asp412/Asp413 (helix α12) seen in the 0° structure, results in 4-8-fold increase in the *K*_m(app)_ value for Gln and a 5-14-fold reduction in the *k*_cat_/*K*_m(app)_ value. Similarly, mutation of Glu213 and Lys376 (helix α11), the interacting pair in the 85° rotated structure led to a 4-fold increase in *K*_m(app)_ value for Gln. The increase in *K*_m(app)_ value for Gln implicates these residues, and the interdomain interaction involving them, in allosteric GATase activation. This apart, we had earlier shown that mutation of Glu374 (helix α11) that interacts with His20 (helix α1) in the 0° structure also impairs GATase activation. The role of α1 in domain crosstalk is supported by NMR experiments on two-subunit type GMPS from *Methanocaldococcus jannaschii* where the binding of substrate-bound ATPPase subunit induced chemical shift changes in residues on helix α1 as well as the loops containing the catalytic triad of the GATase subunit^9^.

Interestingly, although Glu213 is a GATase residue that is far away from the ATPPase active site, the PfGMPS_E213A mutant displays a reduction in the *k*’ for adenyl-XMP formation by 41%. These results validate that rotation of the GATase domain and interactions in the 85° rotated state are integral to allosteric activation of GATase domain as well as catalysis in the ATPPase domain. Corroborating these observations is a study on EcGMPS, in which through juxtaposition of the GATase active site against the ATPPase active site, His186 (analogous to Glu213 in PfGMPS) was seen to form an interdomain salt bridge with Glu383 on α12. Mutation of His186 to Ala resulted in an increase in the *K*_m_ value for Gln by 50-fold and uncoupling of the two reactions^44^.

Taken together, the results from the 0° and 85° interface mutants suggest that the interface helices are conduits for information exchange pertaining to GATase activation and domain crosstalk. In addition, these results suggest that allosteric activation of the GATase domain is a cumulative effect of numerous interdomain interactions in both the 0° and 85° interfaces. It is possible that apart from the two interfaces observed in the available crystal structures, the interdomain interactions in the intermediate states of GATase rotation are also of considerable significance. Our inability to obtain an interface mutant with a very large increase in *K*_m(app)_ value for Gln reflecting GATase activation in the wild type enzyme, suggests the presence of a network of interactions possibly through multiple steps in the trajectory of domain rotation.

### Interface helices link catalysis in ATPPase domain with the GATase activation

The experiments described above have enabled the identification of various residues involved in the functioning of the ATPPase domain. Many of these residues are located on the interface helices α11 and α12 and the lid-loop that is connected to α11. The residues on these structural elements play critical roles in substrate binding (Lys386, Lys411 and Lys415), catalyzing the formation of adenyl-XMP intermediate (Asp371, His388 and His389) and facilitating the reaction of ammonia with adenyl-XMP to generate GMP (Asn390). Whereas residues Asp371, Lys411 and Lys415 on helices α11 and α12 that are essential for ATPPase catalysis have their side chains pointed into the ATPPase pocket, residues Lys376, Glu374 and Asp412/413 have their side chains directed at the interdomain interface where they interact with GATase residues (Figure 7). This arrangement permits the residues on helices α11 and α12 to relay the state of substrate occupancy and/or the formation of adenyl-XMP intermediate to the residues on GATase helix α1. An examination of the structure shows that residues on helix α1 interact with the GATase active site loop which harbours two residues, His208 and Glu210 of the catalytic triad (Figure S2). This permits coupling the events in ATPPase domain (ATP/XMP binding and adenyl-XMP intermediate formation) with GATase activation leading to the co-ordination of the catalytic events across the two domains.

**Figure 7.**
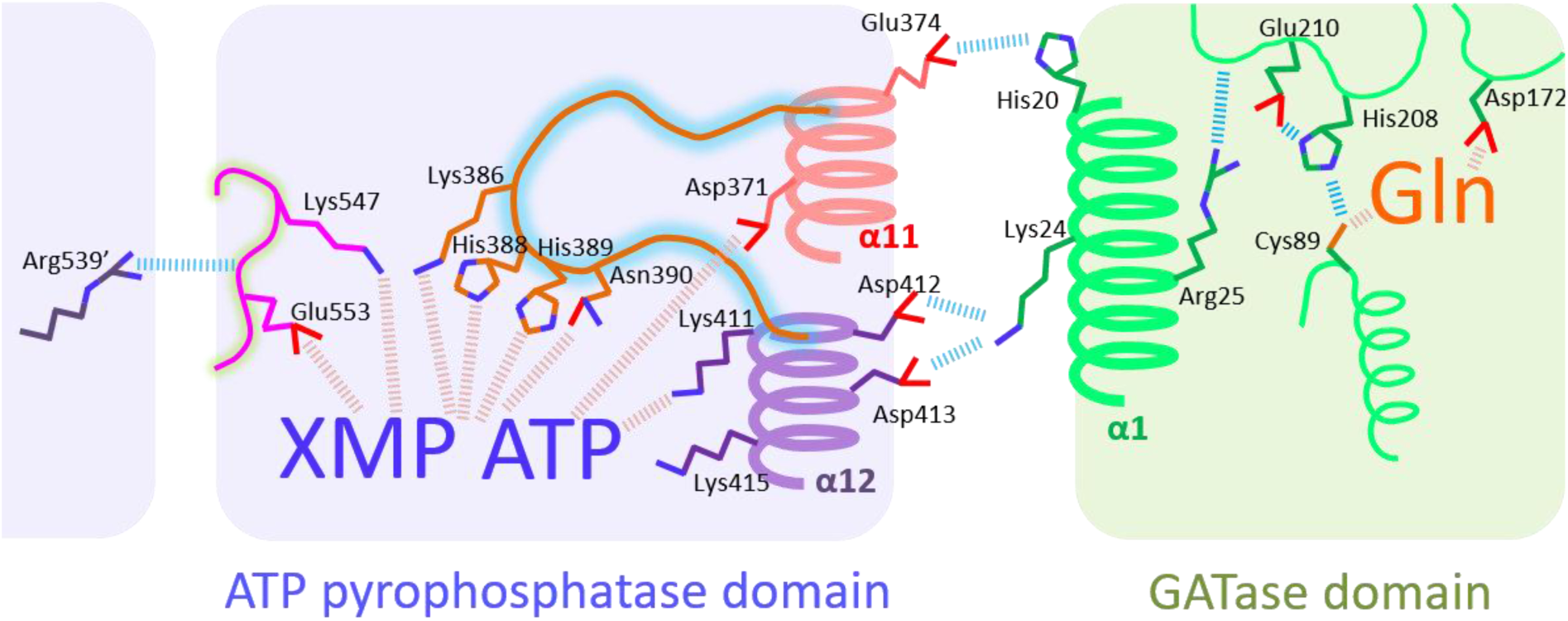
Schematic illustrating the coupling of catalysis in ATPPase domain with allostery. Residues on ATPPase helices α11 and α12 that face the ATPPase pocket along with those on the lid-loop play roles in binding of substrates and ATPPase catalysis. The residues on α11 and α12 that face the interface form salt bridges with residues on GATase helix α1. The interactions in the 0° GATase rotated state is shown here. Residues Lys547 and Glu553 on the C-terminal loop are essential for substrate binding and the conformation of the loop is stabilized by interactions of Arg539’ from the neighbouring chain. Key inter-residue and protein-ligand interactions are shown as blue and brown dashes, respectively. The GATase catalytic triad residues Cys89, His208 and Glu210; and Asp172 which is involved in GATase activation are also shown.

The lid-loop augments these processes since it is connected to the interface helix α11 and thus the conformational dynamics of the lid-loop are linked to GATase rotation. Following substrate binding, the lid-loop closes over the ATPPase active site thereby facilitating the formation of adenyl-XMP intermediate through interactions of His388 and His389. Apart from enabling the synthesis of adenyl-XMP intermediate, the closed conformation of the lid-loop forms a sealed pocket for ammonia tunnelling. This pocket creates a solvent-free environment protecting the adenyl-XMP intermediate from attack by water while also preventing the protonation of ammonia, which would render it incapable of carrying out the nucleophilic attack.

## Conclusions

By mapping various residues that play critical roles in ATPPase catalysis and domain crosstalk, the results presented herein provide insight into the mechanism of synchronization of active sites. The interface helices α11 and α12 play essential roles in ATPPase catalysis, while also serving as conduits for the exchange of information across the two domains. The lid-loop following the interface helix α11 that is linked to the GATase domain, closes over the active site and catalyzes the formation of adenyl-XMP intermediate. Taken together, this study shows the involvement of the flexible lid-loop and the interface helices in synchronizing the activities of the two catalytic pockets. Residues from the neighbouring protomer playing a catalytic role is a feature of PfGMPS that is absent in the human enzyme and this could be exploited for the development of parasite enzyme-specific inhibitors. With the current widespread occurrence of drug resistant malaria, novel drugs targeting essential pathways in the parasite are an imminent requirement.

## Associated Content

### List of supplementary material

Table S1. Sequences of primers used for site-directed mutagenesis.

Table S2. *K*_d_ values for ATP.Mg^2+^ and XMP for the lid-loop mutants.

Table S3. List of interdomain electrostatic interactions in the crystal structures of GMP synthetases where the GATase domain is not rotated (0° rotated structure).

Figure S1. Purification and characterization of PfGMPS WT and mutants.

Figure S2. Interactions of Arg25 on helix α1.

## Author Contributions

SS- conceptualization, structure and sequence analysis, construction of mutants and protein purification, enzyme kinetics, data analysis; NP- construction of mutants and protein purification, enzyme kinetics; LB, SV and NA- solved the structures of PfGMPS used in this study; HB- conceptualization, structure and sequence analyses, data analysis. SS and HB wrote the manuscript with critical inputs from NA. All authors have given approval to the final version of the manuscript.

## Funding Sources

This project was funded by the Department of Biotechnology, Ministry of Science and Technology, Government of India Grants BT/PR11294/BRB/10/1291/2014, BT/PR13760/COE/34/42/2015, and BT/INF/22/SP27679/ 2018; Science and Engineering Research Board, Department of Science and Technology, Government of India Grant EMR/2014/001276; and institutional funding from the Jawaharlal Nehru Centre of Advanced Scientific Research, Department of Science and Technology, India. SS was supported by University Grants Commission junior and senior research fellowships. NP is supported by Indian Council of Medical Research junior research fellowship. HB is a recipient of JC Bose National Fellowship from the Government of India. NA acknowledges funding from the CNRS, the ANR project grant “PLASMOPUR” ANR-17-CE11-0032 and the Fondation Innovation en Infectiologie, “FINOVI project AO12-27”).

## Abbreviations

GATase: glutamine amidotransferase
ATPPase: ATP pyrophosphatase
GMPS: GMP synthetase
PfGMPS: *Plasmodium falciparum* GMP synthetase
WT: wild type
EcGMPS: *Escherichia coli* GMP synthetase
SEM: standard error of the mean

## Notes

The authors declare no competing financial interests.

## Acknowledgements

We acknowledge the constructive criticism offered by Lakshmeesha K. N.

## References

1. Zalkin, H. (1993) The Amidotransferases. Adv. Enzymol. Relat. Areas Mol. Biol. 66, 203–309.

2. Zalkin, H., and Smith, J. L. (1998) Enzymes utilizing glutamine as an amide donor. Adv. Enzymol. Relat. Areas Mol. Biol. 72, 87–144.

3. Massière, F., and Badet-Denisot, M.-A. (1998) The mechanism of glutamine-dependent amidotransferases. Cell. Mol. Life Sci. 54, 205–222.

4. Zalkin, H., and Truitt, C. D. (1977) Characterization of the glutamine site of *Escherichia coli* guanosine 5’ monophosphate synthetase. J. Biol. Chem. 252, 5431–5436.

5. Nakamura, J., Straub, K., Wu, J., and Lou, L. (1995) The glutamine hydrolysis function of human GMP synthetase. Identification of an essential active site cysteine. J. Biol. Chem. 270, 23450–23455.

6. Franco, T. M. A., Rostirolla, D. C., Ducati, R. G., Lorenzini, D. M., Basso, L. A., and Santos, D. S. (2012) Biochemical characterization of recombinant guaA-encoded guanosine monophosphate synthetase (EC 6.3.5.2) from *Mycobacterium tuberculosis* H37Rv strain. Arch. Biochem. Biophys. 517, 1–11.

7. Bhat, J. Y., Venkatachala, R., Singh, K., Gupta, K., Sarma, S. P., and Balaram, H. (2011) Ammonia channeling in *Plasmodium falciparum* GMP synthetase: investigation by NMR spectroscopy and biochemical assays. Biochemistry 50, 3346–3356.

8. Maruoka, S., Horita, S., Lee, W. C., Nagata, K., and Tanokura, M. (2010) Crystal structure of the ATPPase subunit and its substrate-dependent association with the GATase subunit: A novel regulatory mechanism for a two-subunit-type GMP synthetase from *Pyrococcus horikoshii* OT3. J. Mol. Biol. 395, 417–429.

9. Ali, R., Kumar, S., Balaram, H., and Sarma, S. P. (2013) Solution nuclear magnetic resonance structure of the GATase subunit and structural basis of the interaction between GATase and ATPPase subunits in a two-subunit-type GMPS from *Methanocaldococcus jannaschii*. Biochemistry 52, 4308–4323.

10. Jewett, M. W., Lawrence, K. A., Bestor, A., Byram, R., Gherardini, F., and Rosa, P. A. (2009) GuaA and GuaB are essential for *Borrelia burgdorferi* survival in the tick-mouse infection cycle. J. Bacteriol. 191, 6231–6241.

11. Rodriguez-Suarez, R., Xu, D., Veillette, K., Davison, J., Sillaots, S., Kauffman, S., Hu, W., Bowman, J., Martel, N., Trosok, S., Jiang, B., and Roemer, T. (2007) Mechanism-of-action determination of GMP synthase inhibitors and target validation in *Candida albicans* and *Aspergillus fumigatus*. Chem. Biol. 14, 1163–1175.

12. Li, Q., Leija, C., Rijo-Ferreira, F., Chen, J., Cestari, I., Stuart, K., Tu, B. P., and Phillips, M. A. (2015) GMP synthase is essential for viability and infectivity of *Trypanosoma brucei* despite a redundant purine salvage pathway. Mol. Microbiol. 97, 1006–1020.

13. Chitty, J. L., Tatzenko, T. L., Williams, S. J., Koh, Y. Q. A. E., Corfield, E. C., Butler, M. S., Robertson, A. A. B., Cooper, M. A., Kappler, U., Kobe, B., Kobe, B., and Fraser, J. A. (2017) GMP synthase is required for virulence factor production and infection by *Cryptococcus neoformans*. J. Biol. Chem. 292, 3049–3059.

14. McConkey, G. A. (2000) *Plasmodium falciparum*: Isolation and characterisation of a gene encoding protozoan GMP synthase. Exp. Parasitol. 94, 23–32.

15. Bhat, J. Y., Shastri, B. G., and Balaram, H. (2008) Kinetic and biochemical characterization of *Plasmodium falciparum* GMP synthetase. Biochem. J. 409, 263–273.

16. Ballut, L., Violot, S., Shivakumaraswamy, S., Thota, L. P., Sathya, M., Kunala, J., Dijkstra, B. W., Terreux, R., Haser, R., Balaram, H., Balaram, H., and Aghajari, N. (2015) Active site coupling in *Plasmodium falciparum* GMP synthetase is triggered by domain rotation. Nat. Commun. 6.

17. Tesmer, J. J. G., Klem, T. J., Deras, M. L., Davisson, V. J., and Smith, J. L. (1996) The crystal structure of GMP synthetase reveals a novel catalytic triad and is a structural paradigm for two enzyme families. Nat. Struct. Biol. 3, 74–86.

18. Chittur, S. V., Klem, T. J., Shafer, C. M., and Jo Davisson, V. (2001) Mechanism for acivicin inactivation of triad glutamine amidotransferases. Biochemistry 40, 876–887.

19. Raushel, F. M., Thoden, J. B., and Holden, H. M. (2003) Enzymes with molecular tunnels. Acc. Chem. Res. 36, 539–548.

20. Raushel, F. M., Thoden, J. B., and Holden, H. M. (1999) The amidotransferase family of enzymes: Molecular machines for the production and delivery of ammonia. Biochemistry 38, 7891–7899.

21. Mouilleron, S., and Golinelli-Pimpaneau, B. (2007) Conformational changes in ammonia-channeling glutamine amidotransferases. Curr. Opin. Struct. Biol. 17, 653–664.

22. Welin, M., Lehtiö, L., Johansson, A., Flodin, S., Nyman, T., Trésaugues, L., Hammarström, M., Gräslund, S., and Nordlund, P. (2013) Substrate specificity and oligomerization of human GMP synthetase. J. Mol. Biol. 425, 4323–4333.

23. Benson, D. A., Karsch-Mizrachi, I., Clark, K., Lipman, D. J., Ostell, J., and Sayers, E. W. (2012) GenBank. Nucleic Acids Res. 40, D48–53.

24. Uniprot Consortium. (2019) UniProt: A worldwide hub of protein knowledge. Nucleic Acids Res. 47, D506–D515.

25. Katoh Kazutaka, Rozewicki John, Y. K. D. (2019) MAFFT online service: multiple sequence alignment, interactive sequence choice and visualization. Brief. Bioinformatics 20, 1160–1166.

26. Crooks, G. E., Hon, G., Chandonia, J.-M., and Brenner, S. E. (2004) WebLogo: A sequence logo generator. Genome Res. 14, 1188–1190.

27. Berman, H. M., Westbrook, J., Feng, Z., Gilliland, G., Bhat, T. N., Weissig, H., Shindyalov, I. N., and Bourne, P. E. (2000) The Protein Data Bank. Nucleic Acids Res. 28, 235–242.

28. Schrödinger, L. L. C. (2010) The PyMOL Molecular Graphics System. The PyMOL Molecular Graphics System.

29. Barlow, D. J., and Thornton, J. M. (1983) Ion-pairs in proteins. J. Mol. Biol. 168, 867–885.

30. Sarakatsannis, J. N., and Duan, Y. (2005) Statistical characterization of salt bridges in proteins. Proteins Struct. Funct. Genet. 60, 732–739.

31. Guex, N., and Peitsch, M. C. (1997) SWISS-MODEL and the Swiss-PdbViewer: An environment for comparative protein modeling. Electrophoresis 18, 2714–2723.

32. Bradford, M. M. (1976) A rapid and sensitive method for the quantitation of microgram quantities of protein utilizing the principle of protein-dye binding. Anal. Biochem. 72, 248–254.

33. Moyed, H. S., and Magasanik, B. (1957) Enzymes essential for the biosynthesis of nucleic acid guanine; xanthosine 5’-phosphate aminase of *Aerobacter aerogenes*. J. Biol. Chem. 226, 351–363.

34. Fukuyama, T. T. (1966) Formation of an adenyl xanthosine monophosphate intermediate by xanthosine 5’-phosphate aminase and its inhibition by psicofuranine. J. Biol. Chem. 241, 4745– 4749.

35. von der Saal, W., Crysler, C. S., and Villafranca, J. J. (1985) Positional isotope exchange and kinetic experiments with *Escherichia coli* guanosine-5’-monophosphate synthetase. Biochemistry 24, 5343–5350.

36. Vetter, I. R., and Wittinghofer, A. (1999) Nucleoside triphosphate-binding proteins: Different scaffolds to achieve phosphoryl transfer. Q. Rev. Biophys. 32, 1–56.

37. Bork, P., and Koonin, E. V. (1994) A P-loop-like motif in a widespread ATP pyrophosphatase domain: Implications for the evolution of sequence motifs and enzyme activity. Proteins Struct. Funct. Genet. 20, 347–355.

38. Lassila, J. K., Zalatan, J. G., and Herschlag, D. (2011) Biological phosphoryl-transfer reactions: Understanding mechanism and catalysis. Annu. Rev. Biochem. 80, 669–702.

39. Allen, K. N., and Dunaway-Mariano, D. (2016) Catalytic scaffolds for phosphoryl group transfer. Curr. Opin. Struct. Biol. 41, 172–179.

40. Schmelz, S., and Naismith, J. H. (2009) Adenylate-forming enzymes. Curr. Opin. Struct. Biol. 19, 666–671.

41. Yaremchuk, A., Kriklivyi, I., Tukalo, M., and Cusack, S. (2002) Class I tyrosyl-tRNA synthetase has a class II mode of cognate tRNA recognition. EMBO J. 21, 3829–3840.

42. Kobayashi, T., Takimura, T., Sekine, R., Vincent, K., Kamata, K., Sakamoto, K., Nishimura, S., and Yokoyama, S. (2005) Structural snapshots of the KMSKS loop rearrangement for amino acid activation by bacterial tyrosyl-tRNA synthetase. J. Mol. Biol. 346, 105–117.

43. Nakamura, J., and Lou, L. (1995) Biochemical characterization of human GMP synthetase. J. Biol. Chem. 270, 7347–7353.

44. Oliver, J. C., Gudihal, R., Burgner, J. W., Pedley, A. M., Zwierko, A. T., Davisson, V. J., and Linger, R. S. (2014) Conformational changes involving ammonia tunnel formation and allosteric control in GMP synthetase. Arch. Biochem. Biophys. 545, 22–32.

