## Supplementary material for "Helices on interdomain interface couple catalysis in the ATPPase domain with allostery in *Plasmodium falciparum* GMP synthetase"

**List of supplementary material:**

**Table S1.** Sequences of primers used for site-directed mutagenesis.

**Table S2.** $K_d$ values for ATP.$Mg^{2+}$ and XMP for the lid-loop mutants.

**Table S3.** List of interdomain electrostatic interactions in the crystal structures of GMP synthetases where the GATase domain is not rotated (0° rotated structure).

**Figure S1.** Purification and characterization of PfGMPS WT and mutants.

**Figure S2.** Interactions of Arg25 on helix α1.

Table S1. Sequences of primers used for site-directed mutagenesis.

| Mutation | Primer sequence (5'-3') |
|---|---|
| K24L | CTTCCATTTGATTGTATTAAGATTAAACAATATAAAAATATTTAG |
| | CTAAATATTTTTATATTGTTTAATCTTAATACAATCAAATGGAAG |
| R25L | CTTCCATTTGATTGTAAAATTATTAAACAATATAAAAATATTTAGTG |
| | CACTAAATATTTTTATATTGTTTAATAATTTTACAATCAAATGGAAG |
| K160L | GTTTATTTGAAAATATTTTAAGTGACATTACAACGGTTTG |
| | CAAACCGTTGTAATGTCACTTAAAATATTTTCAAATAAAC |
| E213A | CATCCAGAGGTGTATGCATCATTAGATGGAGAATTAATG |
| | CATTAATTCTCCATCTAATGATGCATACACCTCTGGATG |
| K376L | CCAGATATTATTGAAAGTTTATGTTCAAAAAATTTATCAG |
| | CTGATAAATTTTTTGAACATAAACTTTCAATAATATCTGG |
| K386L | CAAAAAATTTATCAGATACTATTTTAACTCATCATAATGTAGG |
| | CCTACATTATGATGAGTTAAAATAGTATCTGATAAATTTTTTG |
| T387A | CAAAAAATTTATCAGATACTATTAAAGCTCATCATAATGTAGGAG |
| | CTCCTACATTATGATGAGCTTTAATAGTATCTGATAAATTTTTTG |
| H388A | CAGATACTATTAAAACTGCTCATAATGTAGGAGGCTTACC |
| | GGTAAGCCTCCTACATTATGAGCAGTTTTAATAGTATCTG |
| H389A | CAGATACTATTAAAACTCATGCTAATGTAGGAGGCTTACC |
| | GGTAAGCCTCCTACATTAGCATGAGTTTTAATAGTATCTG |
| N390A | CAGATACTATTAAAACTCATCATGCTGTAGGAGGCTTACC |
| | GGTAAGCCTCCTACAGCATGATGAGTTTTAATAGTATCTG |
| K411L | GAACCTTTCAAATATTTATTTTTAGATGATGTTAAAACATTATC |
| | GATAATGTTTTAACATCATCTAAAAATAAATATTTGAAAGGTTC |
| D412A | CCTTTCAAATATTTATTTAAAGCTGATGTTAAAACATTATC |
| | GATAATGTTTTAACATCAGCTTTAAATAAATATTTGAAAGG |
| D413A | CCTTTCAAATATTTATTTAAAGATGCTGTTAAAACATTATCTAGAG |
| | CTCTAGATAATGTTTTAACAGCATCTTTAAATAAATATTTGAAAGG |
| K415L | CCTTTCAAATATTTATTTAAAGATGATGTTTTAACATTATCTAGAG |
| | CTCTAGATAATGTTAAAACATCATCTTTAAATAAATATTTGAAAGG |
| R539L | GTGAAGTAAAGGGCGTCAACTTAATATTATATGATGTATCATC |
| | GATGATACATCATATAATATTAAGTTGACGCCCTTTACTTCAC |
| K547L | GAATATTATATGATGTATCATCATTACCACCAGCAACGATTG |
| | CAATCGTTGCTGGTGGTAATGATGATACATCATATAATATTC |
| E553L | CAAAACCACCAGCAACGATTTTATTCGAATGAGAGCTC |

| | |
|---|---|
| | GAGCTCTCATTCGAATAAAATCGTTGCTGGTGGTTTTG |
| E555L | GCAACGATTGAATTCTTATGAGAGCTCCGTCG |
| | CGACGGAGCTCTCATAAGAATTCAATCGTTGC |

Table S2**.** $K_d$ values for ATP.$Mg^{2+}$ and XMP for the lid-loop mutants

| | **$K_d$ (μM)** | |
|---|---|---|
| **Enzyme** | **ATP.$Mg^{2+}$** | **XMP** |
| WT ($K_{m(app)}$) | 119 ± 2 | 7.1 ± 1 |
| H388A | 9.2 ± 1.8 | 0.16 ± 0.03 |
| H389A | 75.6 ± 31.8 | 0.77 ± 0.004 |
| N390A | 2.2 ± 0.2 | 0.04 ± 0.0004 |

The $K_d$ was determined by measuring GATase activity at various concentrations of either ATP.$Mg^{2+}$ or XMP while maintaining the other substrate at fixed saturating concentration which was either 3 mM of ATP.$Mg^{2+}$ or 150 μM of XMP. The glutamate produced was estimated using glutamate dehydrogenase as the coupling enzyme. The mean ± SEM of two experiments is reported.

Table S3. List of interdomain electrostatic interactions in the crystal structures of GMP synthetases where the GATase domain is not rotated (0° rotated structure).

| | Chains | 24 (K/R)(K/R)XRE 28 | 371 DXXES 375 | 411 KD(D/E)V(K/R) 415 |
|---|---|---|---|---|
| ***E. coli*** **(PDB ID: 1GPM)** | A, B, C, D | 25 RRVRE 29 | 340 DVIES 344 | 381 KDEVRK 386 |
| | | Arg25 | | Asp382 [A 2.8, B 3.1, C 2.9, D 5.1] |
| | | Arg28 | | Glu383 [A 2.8, B 2.6, C 2.8, D 2.6] |
| | | Glu29 | | Lys386 [A 5.8, B 5.5, C 5.1, D 5.6] |
| ***C. burnetii*** **(PDB ID: 3TQI)** | A, B, C, D | 25 RRVRE 29 | 339 DVIES 343 | 380 KDEVRK 385 |
| | | Arg25 | | Asp381 [A 3.1, B 3.3, C 3.6, D 2.8] |
| | | Arg28 | | Glu382 [ A 3.0, B 2.8, C 2.7, D 2.9] |
| | | Glu29 | | Lys385 [A 5.6, B 4.4, C 5.8, D 5.7] |
| ***T. thermophilus*** **(PDB ID: 2YWB)** | A | 13 RLIARRLRELR 23 | 320 DVIES 324 | 359 KDEVRE 364 |
| | B, D | 13 RLIARRLRELR 23 | 320 DVIES 324 | 359 KDEVRE 364 |
| | C | 13 RLIARRLRELR 23 | 320 DVIES 324 | 359 KDEVRE 364 |
| | | Arg17 | | Asp360 [A 2.6, B 3.6, C 3.3, D 3.2] |
| | | Arg20 | | Glu361 [B 2.9, C 2.9, D 2.9] |
| ***T. thermophilus*** **(PDB ID: 2YWC)** | A | 13 RLIARRLRELR 23 | 320 DVIES 324 | 359 KDEVRE 364 |
| | B, C, D | 13 RLIARRLRELR 23 | 320 DVIES 324 | 359 KDEVRE 364 |
| | | Arg17 | | Asp360 [A 3.2, B 3.8, C 4.1, D 3.9] |
| | | Arg20 | | Glu361 [A 2.7, B 2.9, C 2.9, D 2.9] |
| ***N. gonorrhoeae*** **(PDB ID: 5TW7)** | A, B, C, D | 17 RLIARRVREAH 27 | 330 DVIES 334 | 371 KDEVRE 376 |
| | | Arg21 | | Asp372 [A 5.2, B 4.9, C 5.5, D 5.0] |
| | | Arg24 | | Glu373 [ A 2.7, B 2.7, C 2.9, D 2.8] |
| | | His27 | | Glu376 [ A 5.4, B 6.0, C 5.5] |
| | | Arg17 | Glu333 [C 3.1] | |
| ***H. sapiens*** **(PDB ID: 2VXO)** | A, B | 39 KVIDRRVRE 47 | 372 DLIES 376 | 415 HKDEVR 420 |
| | | Asp42 | | His415 [A 5.5, B 5.5] |
| | | Arg43 | | Asp417 [B 3.0] |
| | | Arg46 | | Asp417 [A 5.3, B 5.0] |
| | | Arg46 | | Glu418 [A 3.1, B 3.1] |
| ***P. falciparum*** **(PDB ID: 3UOW)** | A | 20 HLIVKRLNNIK 30 | 371 DIIESK 376 | 411 KDDVK 415 |
| | | Lys24 | | Asp413 (5.0) |
| | | Lys24 | | Asp412 (2.7) |
| | B | 20 HLIVKRLNNIK 30 | 371 DIIESK 376 | 411 KDDVK 415 |
| | | His20 | Glu374 (2.9) | |
| | | Lys24 | | Asp413 (4.3) |

| | | | | |
|---|---|---|---|---|
| | | Lys24 | | Asp412 (5.9) |
| | A | 20 HLIVKRLNNIK 30 | 371 DIIESK 376 | 411 KDDVK 415 |
| *P. falciparum* (PDB ID: 4WIM) | | His20 | Glu374 (4.2) | |
| | | Lys24 | | Asp413 (2.8) |
| | B | 20 HLIVKRLNNIK 30 | 371 DIIESK 376 | 411 KDDVK 415 |
| | | Lys24 | | Asp412 (5.3) |

The signature sequence with the numbering corresponding to PfGMPS is mentioned at the top of the table (green). The amino acid sequence of the helices α1, α11 and α12 for every chain analyzed (orange) is mentioned, and the residues corresponding to the signature sequence are underlined. In cases where the side chain of a residue or the entire residue is disordered, the residue is coloured purple. The distance between the salt bridge forming residues in different chains is specified in square brackets. Distances greater than 4.0 Å are highlighted in red.

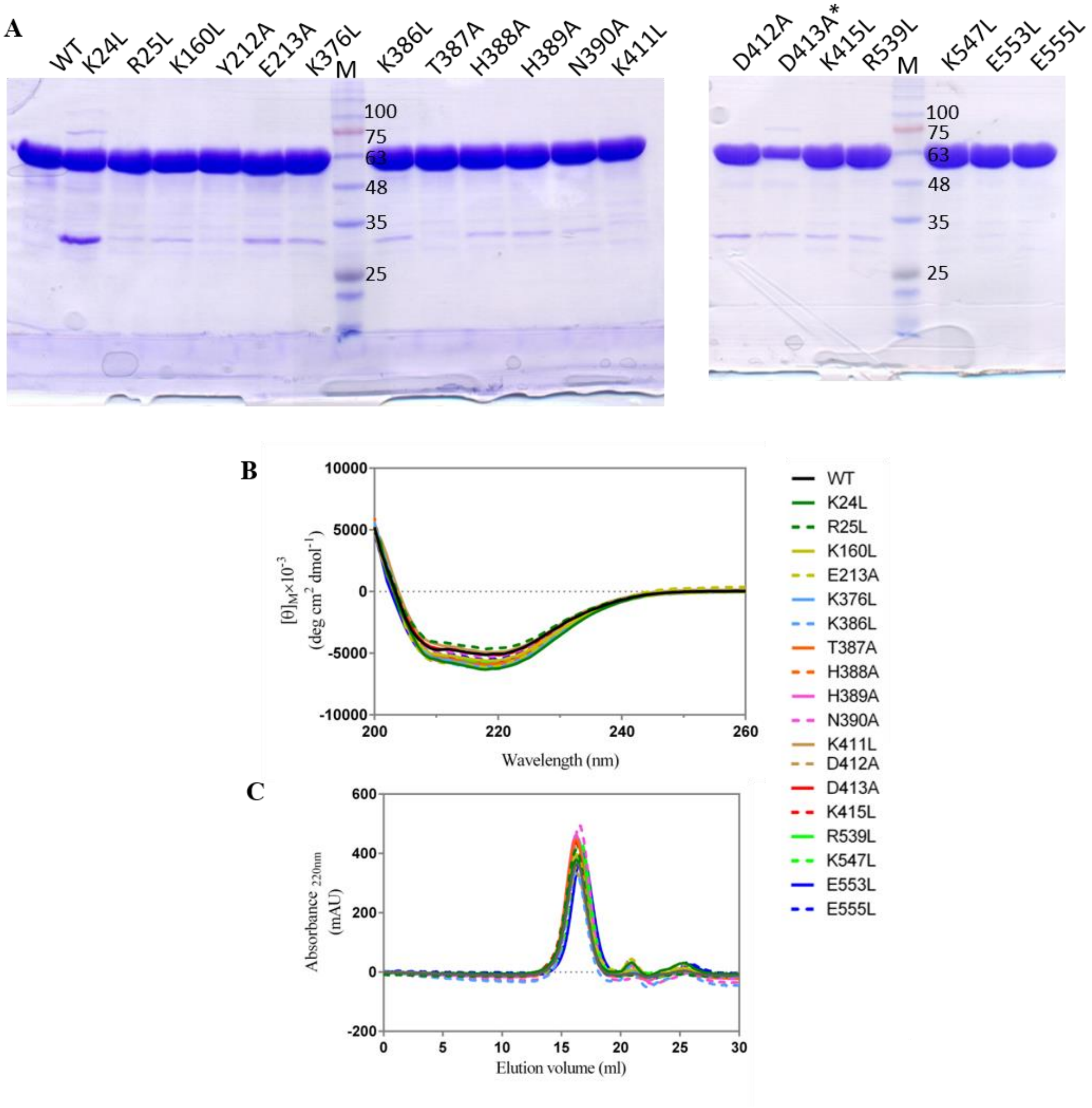


Figure S1. Purification and characterization of PfGMPS WT and mutants. PfGMPS mutant is indicated by the residue, its location in sequence and the residue to which it is mutated. (A) SDS-PAGE of purified proteins. 30 µg of the protein was loaded in all cases except PfGMPS_D413A where 15 µg was loaded. The mutant Y212A is not part of this study. Lane M, molecular mass standards with mass indicated in kDa. (B) The far-UV CD spectra recorded using 5 µM of protein in a buffer containing 6.7 mM Tris-HCl, pH 7.4, 3.3 % (v/v) glycerol, 0.3 mM EDTA and 0.6 mM DTT. (C) Analytical size-exclusion chromatography to determine the oligomeric state. 100 µl of protein (1 mg ml-1) was injected into a column (10 x 300 mm) packed with Sephadex300. The mutants K411L and D413A of PfGMPS were not examined by analytical size-exclusion chromatography.

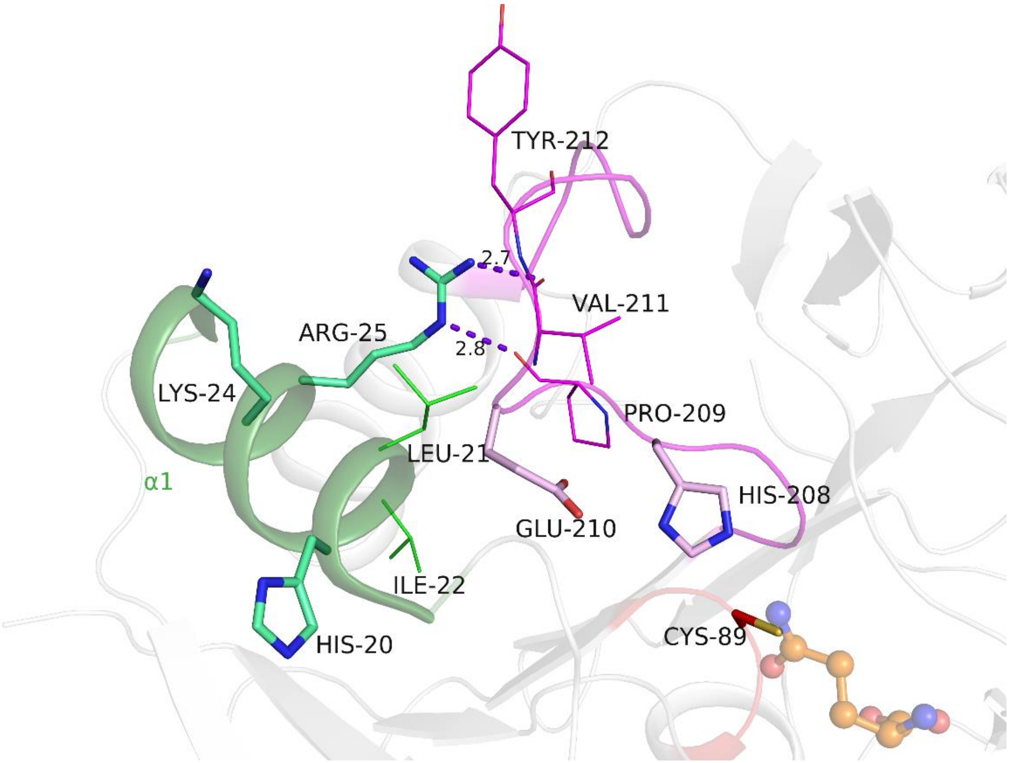


Figure S2. Interactions of Arg25 on interface helix α1 in XMP bound PfGMPS structure (PDB ID 3UOW). The protein backbone is shown as ribbon with α1 coloured green and the loop containing catalytic His208 and Glu210 in pink. The substrate Gln (superposed from 4WIO) is shown in ball and stick representation. Residues on α1 and the loop that are located within 4 Å of each other are shown as lines.
